# Ultrasensitive saliva-based detection of early Alzheimer’s disease biomarkers via nanoparticle-enhanced evanescent scattering microscopy

**DOI:** 10.64898/2025.12.03.692085

**Authors:** Caterina Dallari, Georg Ladurner, Milana Kendrisic, Federica Fenizi Caria, Claudia Manzl, Luisa Ponticelli, Laura Perego, Francesco Goretti, Caterina Credi, Manuela Prokesch, Adelheid Wöhrer, Bernhard Baumann, Rainer Leitgeb, Francesco Saverio Pavone

## Abstract

Alzheimer’s disease (AD) is the most common neurodegenerative disease, yet early diagnosis remains a major challenge. Current cerebrospinal fluid (CSF) assays are invasive and unsuitable for large-scale or repeated screening. Blood-based biomarkers have achieved sensitivities above 90%, but still face challenges of standardization, cost, technical complexity, and the need for sophisticated instrumentation. Saliva offers an attractive, non-invasive solution; however, the low concentrations of AD biomarkers have thus far hindered its clinical applicability. Here, we introduce a saliva-based diagnostic platform that combines Total Internal Reflection Scattering (TIRS) microscopy with antibody-functionalized metallic nanoparticles (NPs), for ultrasensitive, real-time quantification of salivary amyloid-β (Aβ) proteins. Using two established AD mouse models (APP_sl_ and 5xFAD), we found that salivary Aβ₄₂ levels robustly distinguish transgenic from wild-type animals and correlate with brain amyloid deposition. Pooled and stratified analyses suggest these associations are primarily driven by the transgenic-wild-type contrast rather than linear changes within groups. Predictive modeling further confirmed diagnostic utility: in APP_sl_, Logistic Regression and Support Vector Machine (SVM) classifiers both achieved 92% accuracy with balanced sensitivity and specificity, while in 5xFAD, SVM reached 88% accuracy with perfect specificity. These results establish the NPs-enhanced TIRS sensor as a rapid, accurate and non-invasive tool for AD detection via saliva. By addressing a critical unmet need in neurodegenerative disease screening, this platform has strong potential to transform early diagnosis, enable timely interventions, and support biomarker-guided clinical trials. Its simplicity, speed, and scalability, make it well-suited for point-of-care diagnostics and large-scale screening initiatives.

**One Sentence Summary:** Ultrasensitive saliva test detects early Alzheimer’s biomarkers using nanoparticle-enhanced scattering microscopy

## INTRODUCTION

Alzheimer’s disease (AD) is the most prevalent cause of dementia, a progressive neurodegenerative condition marked by a gradual decline in cognitive functions.(1) As of 2020, there were over 55 million people worldwide living with dementia, a number projected to rise to 152 million by 2050, causing increasing stress on healthcare systems in a globally aging population.(2) The economic impact is equally alarming, with global care costs estimated at $2.8 trillion in 2019.(3) In addition to the economic burden, AD imposes psychological, emotional, and physical pressure on patients and their caregivers, often leading to long-term stress, burnout, and reduced quality of life for entire families. Despite decades of research, no definitive cure for AD exists, largely due to its intricate pathogenesis. Only very recently, the first disease-modifying therapies targeting amyloid removal, monoclonal antibodies such as lecanemab and donanemab, have been approved by the FDA and other regulatory agencies worldwide.(4–7) The success of the therapy relies on its ability to modify biochemical pathways and boost immune-mediated clearance of amyloid beta deposits, before irreversible damage in the brain occurs. It thus requires detection before the appearance of the first clinical symptoms.(8) While treatments are most efficient during the early stages of mild cognitive impairment or dementia, they are unlikely to reverse the extensive neuronal damage present during later stages.

To standardize diagnosis, the National Institute of Aging and Alzheimer’s Association (NIA-AA) introduced the A/T/N classification system in 2018. This framework stratifies AD pathology based on biomarkers for amyloid deposition (A), tau pathology (T), and neurodegeneration (N).(9) These biomarkers are typically measured through cerebrospinal fluid (CSF) analysis or neuroimaging techniques, such as positron emission tomography (PET) and structural magnetic resonance imaging (MRI).(10–12) However, CSF-based methods are invasive and unsuitable for broad application or repeated testing due to the procedural complexity and cost of lumbar punctures. Likewise, PET and MRI are limited by high costs, infrastructure requirements, and delayed diagnostic outcomes.(13,14) Since an early and accurate diagnosis is crucial to exploit the window of opportunity between biomarker-driven diagnosis of AD and clinical symptoms, the exploration of more accessible biofluids, including blood,(15) urine,(16) tears,(17) and saliva,(18) has catalyzed attention as a source of in vivo AD biomarkers, even if their use has not yet translated into clinics. While blood-based assays seem promising, their complexity in pre-analytical, cohort-related and analytical/kit-related factors could worsen the reproducibility and sensitivity.(19) Additionally, comorbid pathologies and treatment for other medical conditions affect blood biomarkers for AD.(20,21) In contrast, saliva emerges as an ideal diagnostic medium, as its collection is non-invasive, inexpensive, and does not require hospitalization.

In 2010, Bermejo-Pareja et al. first reported that amyloid-β 42 (Aβ₄₂) levels in saliva were significantly elevated in patients with mild to moderate AD, but not in those with severe AD, compared to cognitively healthy controls, using a commercial enzyme-linked immunosorbent assay (ELISA) kit.(22) In contrast, salivary levels of Aβ₄₀ were unchanged between AD patients and controls. Further studies employing the same ELISA-based approaches corroborated these findings. For instance, Sabbagh et al. and Lee et al. reported a 2.45 and 3-fold increase in Aβ₄₂ concentration in AD patients compared to controls, respectively.(18,23) However, other researchers have encountered challenges in detecting Aβ₄₂ in saliva using either ELISA or Luminex assays, due to the lower amount of salivary proteins in serum or CSF.(23,24) These findings highlight both the potential and the limitations of salivary Aβ₄₂ as a biomarker for AD. While promising, salivary Aβ₄₂ concentrations are likely in the sub-pg/mL or even fg/mL range,(18,22,25) posing a significant challenge for conventional immunoassays. High-sensitivity ELISA kits typically have detection limits in the range of 10-50 pg/mL, which may be insufficient for consistent and accurate quantification of Aβ₄₂ in saliva, thereby limiting their applicability in clinical settings.(26,27) Therefore, there is an urgent need for highly sensitive alternative methods that can detect ultralow concentrations of biomarkers through saliva screening, to facilitate the identification of individuals at higher risk of AD exploiting safe and cost-effective procedures.

To this aim, we propose a novel diagnostic sensor based on Total Internal Reflection Scattering (TIRS). In TIRS, the reduced penetration depth of the excitation field is exploited to limit the investigated volume to a small portion of the sample, thus reducing the background contribution from deeper regions.(28–31) Only molecules located within the excited region will be illuminated, enhancing the contrast and improving the detection sensitivity.(32) Moreover, metallic nanoparticles (NPs) with intrinsically large scattering crosssections can be functionalized to specifically self-assemble onto AD biomarkers of interest, further increasing the sensitivity of the analysis and enabling the detection of biomarkers at ultralow concentrations (down to the attogram per milliliter range, i.e., 6x better than conventional ELISA assays). This breakthrough in sensitivity has the potential to unlock the analysis of biomarkers in less invasive, more accessible biological fluids such as saliva, which until now have remained underutilized for AD diagnostics due to the extremely low concentration of target analytes, marking a significant step forward from the current reliance on CSF analysis and neuroimaging.

Here, we apply this NPs-enhanced TIRS platform to two established AD mouse models (human APP_sl_ and 5xFAD),(33) and demonstrate that salivary Aβ₄₂ levels reliably discriminate transgenic from wild-type animals and exhibit significant correlations with brain Aβ₄₂ concentrations when all animals (wt and tg) are analyzed together, with the strongest associations observed in the hippocampus. Predictive modeling using machine learning approaches further confirms its diagnostic potential, with accuracies exceeding 88-92% depending on the model. Importantly, the platform is rapid, scalable, and non-invasive, addressing a critical unmet need in neurodegenerative disease screening. By enabling ultrasensitive detection of salivary biomarkers, this approach holds promise for translation into point-of-care diagnostics and large-scale screening initiatives, ultimately facilitating earlier intervention and biomarker-guided clinical trials.

## Results

The reported compact system using prism-based total internal reflection (TIR) has been specifically adapted for the ultrasensitive detection of scattering signals from AD biomarkers.(34) The working principle of the setup is detailed in Methods and graphically reported in Figure S1 and S2. The shallow penetration depth of the evanescent-wave field generated by TIR allows selective excitation of analytes near the sensor surface, effectively reducing background signals from deeper regions of the solution (Figure 1a). Moreover, to enhance detection sensitivity, metallic nanoparticles (NPs), and in particular gold NPs with high scattering cross-sections, are functionalized to specifically bind the biomarkers of interest.(35–38) In this study, NPs were functionalized to selectively capture AD biomarkers (Aβ₄₂ and Aβ₄₀).(39) To validate the performance of the optical system to target AD biomarkers, we implemented a protocol for analyzing biological samples containing the target biomarkers (Figure 1b). Briefly, biological fluids were collected and incubated on a sensing surface functionalized with capture antibodies specific to β-amyloid peptides. After the target peptides were immobilized, gold NPs conjugated with detection antibodies (NPs-Ab) were introduced. These NPs-Ab bind selectively to the captured biomarkers, forming a sandwich immunoassay (Supplementary Figures S3 and S4). Thanks to the large scattering cross-section of gold NPs, their binding can be sensitively detected using the TIRS setup. The interaction between the EW and the NPs-Ab complexes generates a strong scattering signal, enabling rapid biomarker detection within minutes. Upon binding to target peptides, the nanoparticles self-assemble, leading to measurable changes in elastic scattering intensity. The concentration of Aβ peptides is calculated from the density of nanoparticles, which directly correlates with peptide abundance. To quantitatively evaluate the performance of the TIRS setup to detect AD biomarkers highly diluted in liquid samples, scattering measurements were performed using known concentrations of synthetic Aβ₄₀ and Aβ₄₂ in solution (Figure 1 c,d). We observed a strong relationship between the calculated NPs density change and the biomarkers’ logarithmic concentration in the range of 10^0^ to 10^-9^ ng/mL. The coefficients of determination of the regression line (R^2^) were 0.997 and 0.956 for Aβ₄₂ and Aβ₄₀, respectively (Supplementary Figure S5). We further examined the setup capability to specifically recognize different amyloid species-namely, Aβ₄₂ and Aβ₄₀-when both were present in solution. No significant changes in NPs density were observed when measuring the targeted amyloid in solutions containing one biomarker or both, successfully confirming that our method can distinguish between the two Aβ species, respectively, without the interference of other non-target Aβ species down to the attogram/mL (10^-18^ g/mL) range (Figure 1e, p-values reported in table S1 in Supporting Info file). These findings imply that the TIRS sensing platform can selectively recognize AD biomarkers without significant interference from other non-target proteins.

**Figure 1:**
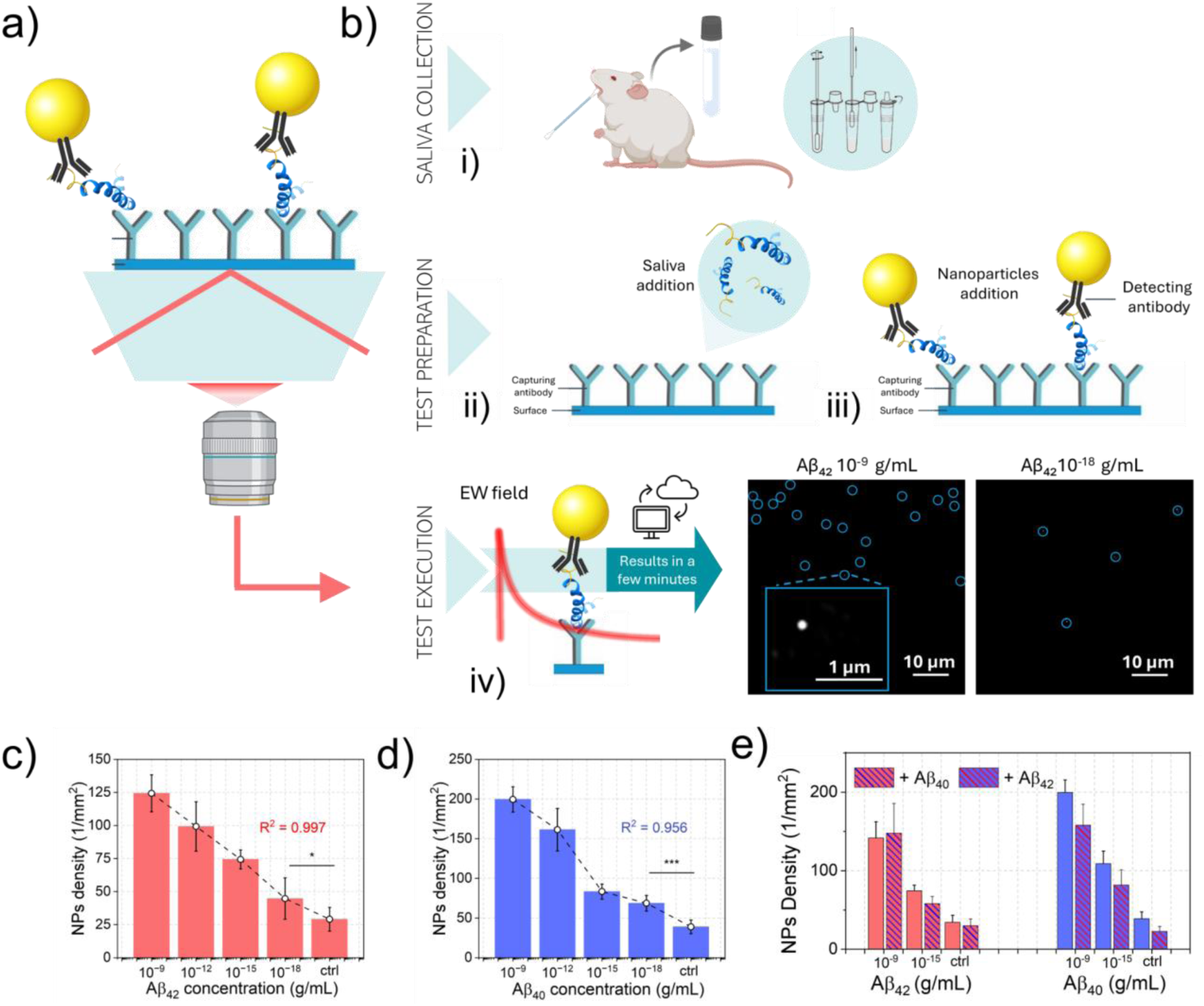
Nanoparticle-enhanced TIRS assay for Aβ biomarker detection. (a) Schematic representation of total internal reflection scattering (TIRS)-based optical setup. (b) Workflow of the assay: (i) non-invasive saliva collection from mice using a cotton swab; (ii) incubation on antibody-functionalized substrates; (iii) binding of NPs conjugated with ad-hoc antibodies (NPs-Ab) to self-assemble on target peptides; and (iv) readout of scattering signals, exploiting the EW field as an excitation source. (c,d) Scattering measurements were performed with synthetic beta amyloids 42 and 40 (Aβ₄₂ and Aβ₄₀) concentrations, ranging from 10^-9^ to 10^-18^ g/mL for each biomarker; n>30. Error bars indicate the standard error. Mean NPs density as a function of Aβ₄₂ (c) and Aβ₄₀ (d) concentration was graphed both in histogram and dot forms. (e) Selectivity of the TIRS-based sensor toward individual AD biomarkers and their mixture was tested. The concentrations Aβ₄₂ and Aβ₄₀ were 0, 10^-9^ and 10^-15^ g/mL. n>30 and error bars indicate the standard error. R^2^ is the coefficient of determination and quantifies how well the regression model explains the variability of the observed data.

We then investigated the applicability of the TIRS-based sensing platform using samples collected from transgenic AD mouse models (tg) and wild-type (wt) controls in both 5xFAD and APP_sl_ mouse lines. Detailed demographic characteristics of the animals are provided in Supplementary Table S2. The same protocol described for synthetic amyloid and graphically reported in Figure 1b was implemented. Saliva samples were diluted 1:1 with Phosphate Buffer Saline (PBS) buffer and incubated on the sensing surface functionalized *ad-hoc* to recognize amyloid peptides. Scattering signals from NPs-Ab complexes assembled on peptides were then collected. Statistical analyses revealed significant differences in biomarker concentrations between wt and tg groups in both mouse models. Repeated measurements collapsed to a single subject-level value using the within-animal median for each biomarker (when only one measurement was available, that value was retained). All subsequent analyses are based on these medians. In the APP_sl_ model, average Aβ₄₂ levels were approximately 2.5 times higher in tg mice compared to wt controls (167 vs. 67; Figure 2 a,i and a,ii). Similarly, in the 5xFAD model, the increase was approximately 3.3-fold (231 vs. 71; Figure 2 e,i and e,ii). The differences were highly significant (p < 0.0001, unpaired t-test) and align with recent literature reporting elevated amyloid peptide levels in saliva samples from diseased subjects.(18,22,23) Instead, average Aβ₄₀ levels were 2.3 times higher in tg mice compared to wt in the APP_sl_ model (150 vs. 65; Figure 2b,i and b,ii), and 1.91 times higher in the 5xFAD model (188 vs. 98; Figure 2f,i and f,ii), with p-values slightly above 0.01 in both models. The average Aβ₄₂/Aβ₄₀ ratio in APP_sl_ tg mice was higher on average (1.43 vs. 0.9; Figure 2 c,i and c,ii), corresponding to an approximately 1.5fold increase, but this difference did not reach statistical significance (p = 0.26). In contrast, the 5xFAD tg mice exhibited a statistically significant 2.38-fold increase in the average Aβ₄₂/Aβ₄₀ ratio compared to wt controls (1.19 vs. 0.5; p < 0.01; Figure 2 g,i and g,ii). All data, including mean values, differences and statistics are reported in Supplementary Table S4. Also, based on the coefficients of variation (CV), the relative dispersion appears comparable between groups. Receiver operating characteristic (ROC) curve analyses confirmed the high discriminative power of Aβ₄₂, with area under the curve (AUC) values of 0.98 in both APP_sl_ and 5xFAD models. In contrast, Aβ₄₀ and Aβ₄₂/Aβ₄₀ ratio showed weaker discriminative performance. These findings highlight Aβ₄₂ as the most robust and reliable biomarker for distinguishing AD transgenic from control animals in both AD models. Based on these results, subsequent analyses were focused on Aβ₄₂ quantification.

**Figure 2:**
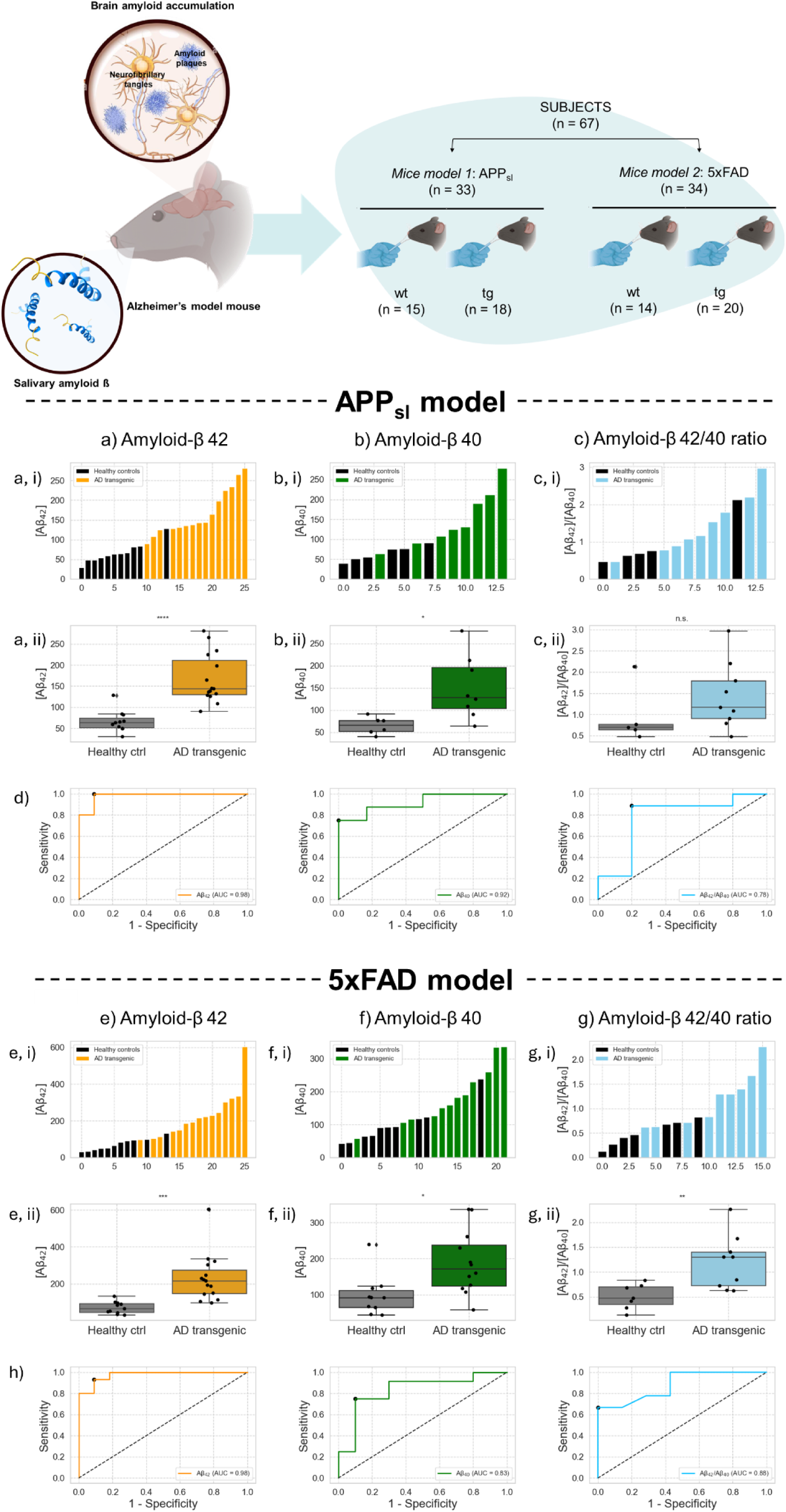
Salivary Aβ biomarker quantification in APP_sl_ (n=33) and 5xFAD mice (n=34). (a-c) Waterfall plots and box plots for APP_sl_ animals showing the calculated levels of Aβ₄₂ (a i,ii), Aβ₄₀ (b i,ii), and the Aβ₄₂/Aβ₄₀ ratio (c i,ii) in saliva of AD mice (n=18) and mouse controls (n=15). X-axes are expressed in terms of NPs density. (d) Receiver operating characteristic (ROC) curves of the TIRS sensor for Aβ₄₂, Aβ₄₀, and the Aβ₄₂/Aβ₄₀ ratio were used as a predictor, respectively. (e-g) Waterfall plots and box plots for 5xFAD animals showing the calculated levels of Aβ₄₂ (e i,ii), Aβ₄₀ (f i,ii), and the Aβ₄₂/Aβ₄₀ ratio (g i,ii) in saliva of AD mice (n=20) and mouse controls (n=14). X-axes are expressed in terms of NPs density. (h) ROC curves of the TIRS sensor for Aβ₄₂, Aβ₄₀, and the Aβ₄₂/Aβ₄₀ ratio were used as a predictor, respectively. Data are shown as boxplots, where the central line represents the median, the box indicates the interquartile range (IQR), and the whiskers extend to 1.5 times the IQR. Outliers correspond to the individual points that exceed the whiskers. Median: represents the central value between the minimum and maximum. Interquartile Distance (IQD): the distance between the Upper and the Lower Quartiles. Max: maximum value. Min: minimum value. Statistical analysis was performed using a two-sample unpaired Student’s t-test with non-statistical significance (NS) >0.05; *p<0.05; **p <0.01; ***p<0.001.

### Biochemistry and histology

Enzyme-linked immunosorbent (ELISA) assays were employed to quantify amyloid levels in the hippocampus and cortex of the left hemisphere in both 5xFAD and APP_sl_ mouse models. In both transgenic lines, amyloid concentrations were elevated relative to wild-type controls for both Aβ species, with a minimum 5.7-fold increase, measured between wt to tg 5xFAD mice for Aβ₄₂ in the cortex, and a maximum 550-fold increase for Aβ₄₀ values in the hippocampus of APP_sl_ mice. Measured values for both amyloid species, brain parts and models can be found in the Table in Figure 3i. In APP_sl_ mice, amyloid levels were substantially higher in the hippocampus compared to the cortex, suggesting region-specific differences. (Figure 3c,d) In 5xFAD mice, Aβ₄₀ levels were considerably elevated in the hippocampus relative to the cortex (Figure 3a), whereas Aβ₄₂ levels were present at comparable concentrations in both regions, with a slight increase observed in the cortex. The right hemibrains of animals were used for histological staining with Aβ₄₀ and Aβ₄₂ for both mouse lines. For the 5xFAD mice, small core plaques and cerebral amyloid angiopathy (CAA) were identified for both staining. Aβ₄₀ was found exclusively in the hippocampus (Figure 3e). Aβ₄₂-stained plaques appeared predominantly in the hippocampus, but also in the cortex and midbrain (Figure 3f). For tg APP_sl_ mice, Aβ₄₀ (Figure 3g) and Aβ₄₂ plaques (Figure 3h) appeared diffusely in the hippocampus and cortex, in the form of large confluent plaques with corona and CAA. Plaques were nearly absent from the midbrain, with rare exceptions. Non-transgenic mice of both lines did not exhibit any plaques. The stains were entirely negative.

**Figure 3:**
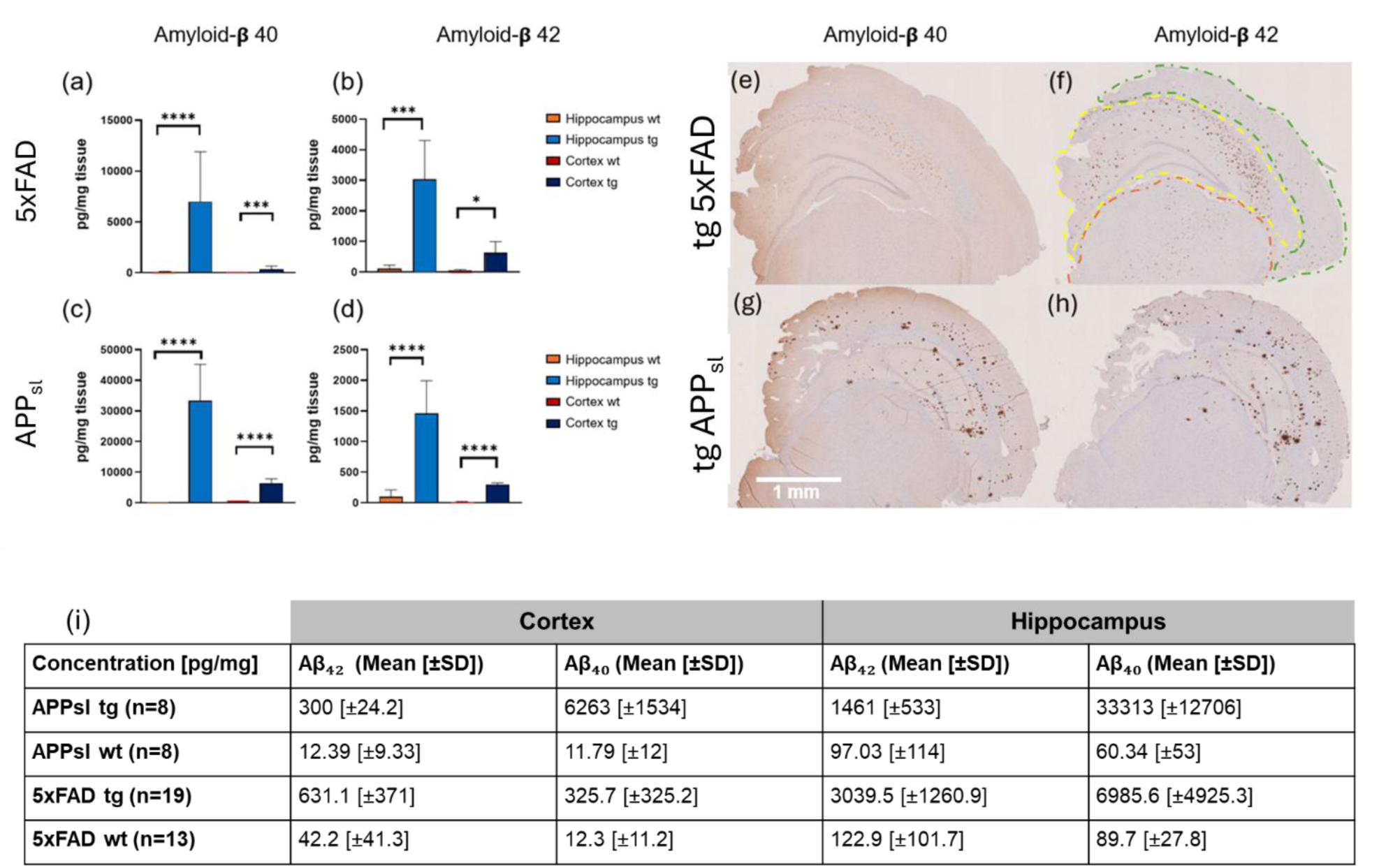
Brain amyloid quantification in APP_sl_ and 5xFAD mice. (a,b) ELISA-based Aβ₄₀ and Aβ₄₂ concentrations in hippocampus and cortex of tg and wt mice for 5xFAD (n(tg) = 19, n(wt) = 13). (c,d) Equivalent analysis in APP_sl_ mice (n(tg) = 8, n(wt) = 8). Mean concentration of amyloid expression measured by ELISA is depicted as bar graphs ± standard deviation (SD) of xy-to-xy mice in panels (a-d). (e-h) Histological staining of right hemibrain sections of investigated mouse models showing parts of the hippocampus, cortex and midbrain, outlined in yellow, green and orange panel (f), respectively. (e) Aβ₄₀ staining of a 5xFAD transgenic mouse. (f) Aβ₄₂ staining of a 5xFAD mouse. (g) Aβ₄₀ staining of an APP_sl_ mouse. (h) Aβ₄₂ staining of an APP_sl_ mouse. (i) Summary of mean Aβ concentrations from ELISA. (*p<0.05, ***p<0.001, ****p<0.0001, determined with the unpaired t-test).

### Correlation between amyloid measurements in saliva and biochemistry

To investigate the relationship between salivary and brain levels of Aβ₄₂, a correlation analysis was conducted between Aβ₄₂ concentrations in saliva, hippocampus and cortex. The analysis was conducted on 15 animals for APP_sl_ for both saliva-hippocampus and saliva-cortex. For 5xFAD, 24 animals provided observation pairs, with n=23 for saliva-hippocampus and n=24 for saliva-cortex. As reported in Figure 4a, in APP_sl_ correlation pooled across wt and tg animals, salivary Aβ₄₂ showed a strong positive correlation with hippocampal Aβ₄₂ (r = 0.901, p < 0.001; ρ = 0.837, p < 0.001) and with cortical Aβ₄₂ (r = 0.727, p = 0.002; ρ = 0.649, p = 0.009). Hippocampal and cortical levels were also closely correlated (r = 0.843, p < 0.001; ρ = 0.664, p = 0.007). Similarly, in 5xFAD (Figure 4b), salivary Aβ₄₂ correlated positively with hippocampal Aβ₄₂ (Pearson r = 0.610, p = 0.002; Spearman’s ρ = 0.609, p = 0.002). The saliva–cortex association was weaker by Pearson (r = 0.264, p = 0.212) but significant by Spearman (ρ = 0.637, p < 0.001), suggesting a monotonic relationship influenced by non-linearity or outliers. Hippocampal and cortical Aβ₄₂ were also associated (r = 0.543, p = 0.002; ρ = 0.687, p < 0.001). Overall, pooled analyses indicate that salivary Aβ₄₂ tracks brain Aβ₄₂, particularly hippocampal levels, in both models, with stronger and more consistent evidence in APP_sl_. When stratified by genotype, in APP_sl_-tg the saliva-hippocampus correlation remained significant (Pearson r = 0.795, p = 0.018; Spearman’s ρ = 0.714, p = 0.047), whereas other within-group pairs were not significant. In 5xFAD, within-group analyses did not reveal significant correlations, except for the very high hippocampus-cortex association in wt animals (Pearson r = 0.993, p < 0.001). These findings indicate that the positive saliva-hippocampus association observed in pooled data is largely driven by the difference between tg and wt groups rather than by linear trends within groups (a Simpson’s paradox situation). Stratified analyses should be interpreted with caution, as the reduced sample sizes yield fewer stable estimates. Violin plots in Figure 4c and d represent the distribution of values using a Gaussian kernel density estimate, providing a smoothed view of the data in addition to the median and interquartile range. They reveal clear increases in Aβ₄₂ concentrations in tg animals compared to wt across saliva, hippocampus, and cortex. In both APP_sl_ and 5xFAD, tg animals exhibited higher medians, but variability was substantially greater in 5xFAD, as reflected by the broader distributions. Wt animals, by contrast, clustered tightly at low values. Thus, both models highlight a shift in central tendency between genotypes, but variability is markedly higher in 5xFAD.Regression plots in Figure 4e and f provide a graphical illustration of these associations, showing that apparent positive trends in pooled data reflect the separation between tg and wt animals, while within-group slopes are weak and highly uncertain, as evidenced by the wide 95% confidence intervals (shaded areas) and the limited number of observations. Full regression results for pooled analyses are reported in Supplementary Figure S6.

**Figure 4:**
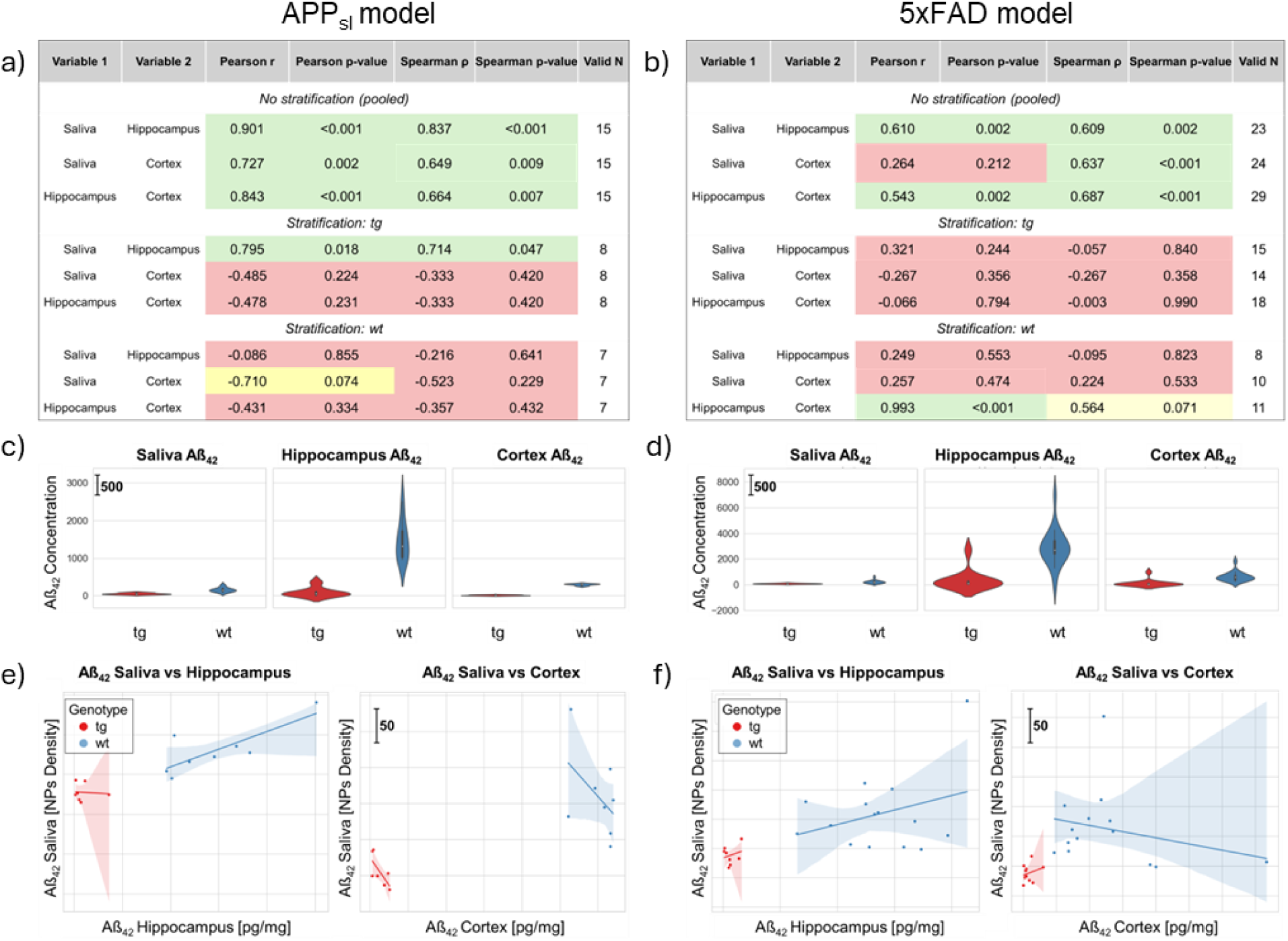
Comparative analysis of salivary and brain Aβ₄₂ concentrations. (a,b) Correlation between Aβ₄₂ levels in saliva, hippocampus, and cortex, in APP_sl_ (a) and 5xFAD mice (b), assessed by Pearson’s r and Spearman’s ρ. (c,d) Distribution of Aβ₄₂ concentrations in saliva, hippocampus and cortex stratified by genotype for APP_sl_ (c) and 5xFAD (d). (e,f) Regression analyses to compare Aβ₄₂ levels in saliva versus hippocampus (left panel) and in saliva versus cortex (right panel), with shaded bands indicating 95% confidence intervals for APP_sl_ (e; n(tg)=8; n(wt)=7) and for 5xFAD (f; n(tg)=15; n(wt)=8 saliva-hippocampus; tg n(tg)=14; n(wt)=10 saliva-cortex).

### Predictive models-Support Vector Analysis

To assess the diagnostic potential of salivary Aβ₄₂ concentration, we trained Logistic Regression and Support Vector Machine (SVM) classifiers with Leave-One-Out Cross-Validation (LOOCV). Uncertainty in performance estimates was quantified by bootstrap resampling (B = 1000), which provided 95% confidence intervals (CI) for AUC, accuracy, and optimal decision threshold expressed in terms of salivary Aβ₄₂ concentration. In the APP_sl_ model, Logistic Regression achieved an AUC of 0.95 (95% CI: 0.84-1.00) with an accuracy of 0.92 (95% CI: 0.85-1.00) and an F1-score of 0.93. The Youden-optimal threshold corresponded to a salivary Aβ₄₂ concentration of 104 pg/mL (95% CI: 83.9-134.5). Performance metrics indicated balanced sensitivity (93%) and specificity (91%), with misclassified cases clustering near the decision threshold, reflecting the inherent difficulty in classifying borderline biomarker levels. The SVM classifier yielded nearly identical results (AUC 0.95, accuracy 0.92, F1-score 0.93), with an optimal threshold of 104 pg/mL, a slightly wider confidence interval (95% CI: 81.9-135). The hyperparameter search consistently selected a linear kernel, suggesting that with the current dataset size, a linear decision boundary was sufficient and that no gain was obtained from more flexible kernels. In 5xFAD model, Logistic Regression achieved an AUC of 0.93 (95% CI: 0.83-1.00), with an accuracy of 0.86 (95% CI: 0.77-1.00) and an F1-score of 0.85. The Youden-optimal threshold corresponded to a salivary Aβ₄₂ concentration of 143 pg/mL (95% CI: 89-186). Performance metrics highlighted perfect specificity (100%) but lower sensitivity (73%), indicating that misclassifications were primarily false negatives, i.e. transgenic mice with biomarker values close to the threshold incorrectly classified as wt. The SVM classifier showed very similar results, with an AUC of 0.95 (95% CI: 0.82-1.00), accuracy of 0.88 (95% CI: 0.77-1.00), and F1-score of 0.89. The optimal decision threshold was slightly lower than in Logistic Regression (138 pg/mL, 95% CI: 84-193), but the operating characteristics remained unchanged: high specificity was preserved, while sensitivity improved only marginally to 80%. As in the Logistic model, errors clustered around the decision boundary, reflecting the difficulty of classifying borderline cases. Overall, Logistic Regression and SVM produced nearly identical results within each model, reflecting the adequacy of linear decision functions for the available data (Logistic Regression results reported in Supplementary Figure S7). More flexible kernels did not yield systematic improvements, likely due to the limited sample size and the predominantly linear separation between wt and tg animals. Bootstrap confidence intervals provide further insight into the robustness of these results (Figure 5f). In both models, accuracy and AUC estimates were consistently high, with 95% confidence intervals spanning ranges that remained within good to excellent performance (e.g., AUC 0.83–1.00). Confidence intervals for the optimal salivary Aβ₄₂ thresholds were broad (e.g., 84–193 pg/mL in 5xFAD), indicating that while the biomarker reliably discriminates between genotypes, the precise cut-off cannot yet be determined with high precision. Thus, larger cohorts will be required to refine threshold estimates and narrow the range of performance metrics but salivary Aβ₄₂ emerges as a robust and reproducible marker of transgenic status.

**Figure 5:**
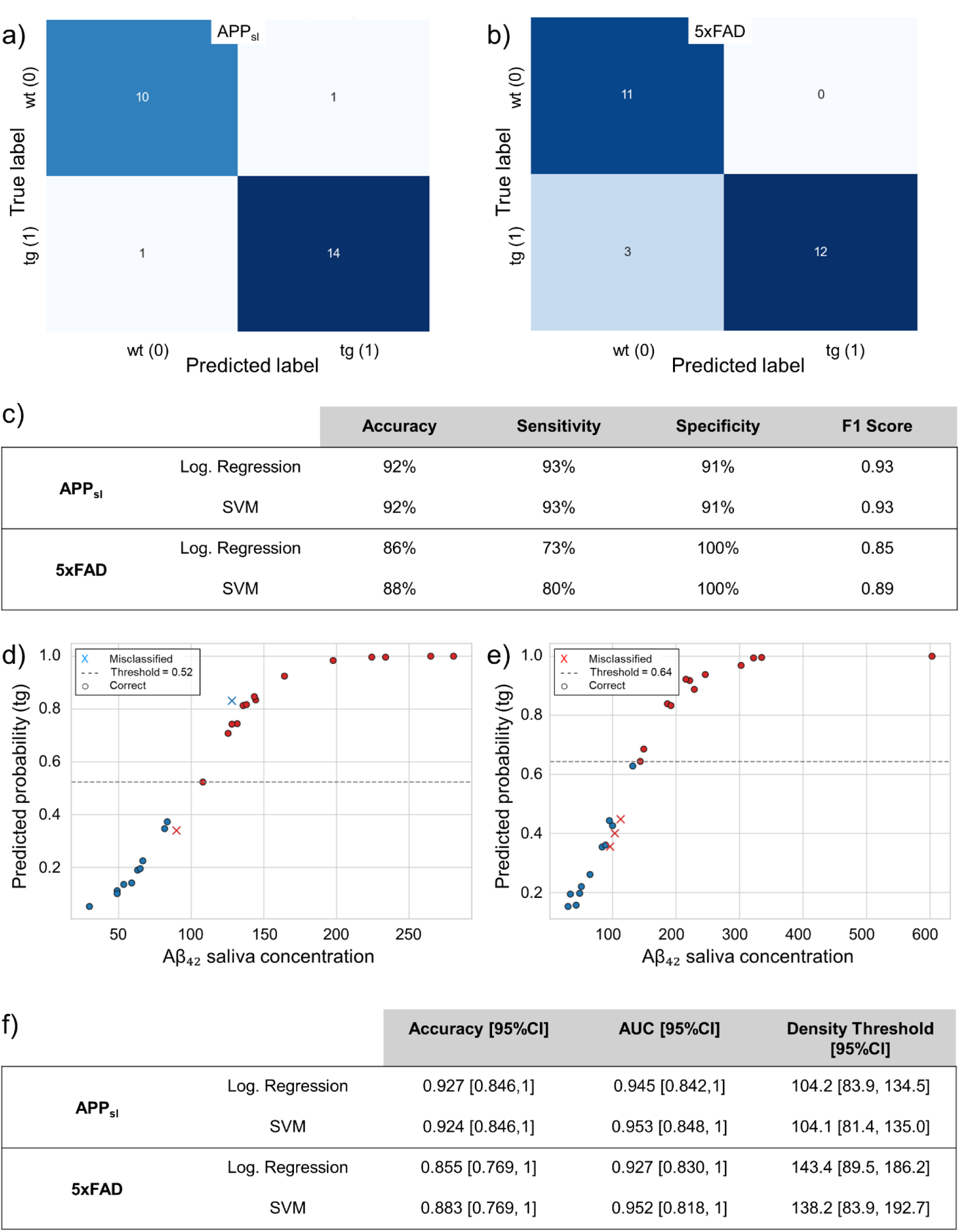
Predictive modeling of salivary Aβ₄₂ for AD detection. (a,b) Confusion matrices of support vector machine (SVM) classifier in APP_sl_ and 5xFAD models based on salivary Aβ₄₂ concentrations. The SVM classifier was evaluated using LOOCV. (c) Summary of classification performance metrics (accuracy, sensitivity, specificity and F1-score) for Logistic Regression and SVM. (d,e) Probability plots of predicted transgenic status based on salivary Aβ₄₂. (f) Summary of accuracies, AUC, and optimal density thresholds with 95% bootstrap confidence intervals. For each metric, the first value indicates the point estimate, while the values in brackets denote the 2.5th and 97.5th percentiles of the bootstrap distribution.

## Discussion

Total internal reflection scattering (TIRS)-based optical detection, coupled with antibody-functionalized gold nanoparticles, demonstrates superior sensitivity and diagnostic precision compared with conventional sandwich-type enzyme-linked immunosorbent assays (ELISA). Our system achieves ∼6-fold lower detection limits than commercially available ELISAs,(26,27) enabling the quantification of Aβ biomarkers in saliva at concentrations previously considered unattainable. While alternative approaches, such as surface plasmon resonance (SPR), fluorescence-based assays, and electrochemical sensors, have been explored for salivary AD biomarker detection, they often face limitations in sensitivity, surface preparation, or scalability for point-of-care applications. A 2020 study explored the use of ELISA to detect AD biomarkers in the saliva of a small patient cohort, reporting a 2.5-fold increase of Aβ₄₂ for AD patients and proposing saliva as a potential approach for AD diagnosis.(18) Our observed 2.5-to 3-fold increase of average Aβ₄₂ levels in tg animals, mirrors these earlier human findings. However, ELISA-based protein detection poses critical downsides for the application in the clinical settings. Lower sensitivity in addition to high cost is a crucial downside of ELISA assays. Additionally, the inflexibility of the immunoassays, allowing for testing of only one marker per kit, is a critical downside of ELISA, compared to the TIRS platform presented in this study. TIRS can detect multiple biomarkers in one sample, making the testing of multiple markers in one measurement feasible, greatly reducing cost and time needed. The state of the art for AD biomarker testing includes blood tests and CSF sampling. Several studies were conducted to determine the feasibility of blood-based biomarkers and the accuracy of the tests applied. In 2024 a study conducted by Palmqvist et al., reported an accuracy of 90% for the combination of the APS2 test for Aβ₄₀/Aβ₄₂ ratio and p-tau217 or the percentage of p-tau217 alone for AD diagnosis in patients in primary care.(40) The FDA-approved Lumipulse G β-amyloid ratio (42/40) blood test reports a sensitivity of 91.7% and a specificity of 97.3%.(41) In a study by Barthélemy et al. from 2024, a blood test for plasma p-tau217 determined Aβ status with an accuracy of 89-90%, increased to 95% with a two-cutoff strategy, in more than 80% of confidence interval patients using liquid chromatography-tandem high resolution mass spectroscopy analysis, outperforming standard CSF tests.(42) It should, however, be emphasized that neither liquid chromatography nor mass spectroscopy are inexpensive or commonly available analytical tools and might not qualify as testing devices for broad screening. In comparison, to blood and CSF tests, our proof-of-concept study achieved average sensitivities of 88-92% for salivary Aβ₄₂ detection. Machinelearning-based analyses achieved accuracies of 88-92% in distinguishing wild-type from transgenic mice, positioning the diagnostic capabilities of the TIRS platform on par with up-to-date blood assays, while offering advantages of non-invasiveness, simplicity and scalability. Including additional Aβ and p-tau in preclinical and clinical settings, as well as overall refinement of the method, will likely further increase the sensitivity. Additionally, salivary Aβ₄₂ concentrations were associated with brain amyloid levels in both mouse models, with the strongest and most consistent relationships observed with hippocampal deposition. In APP_sl_, these associations were evident in pooled analyses and for saliva–hippocampus, also persisted within transgenic animals. In 5xFAD, saliva–hippocampus correlations were significant at the group level but not within genotypes, indicating that the observed associations are largely driven by the transgenic versus wild-type contrast rather than by linear trends within groups. Since AD pathology typically begins in the hippocampus and later spreads to the cortex in both humans(43) and the 5xFAD model(44), we attribute this discrepancy to the age difference between the two cohorts. The investigated APP_sl_ mice, aged 16-17 months, were nearly twice as old as 5xFAD mice (8.5–9.5 months), providing a plausible explanation for why cortical correlations are detectable only in the former. In APP_sl_ mice, disease progression had advanced sufficiently to involve the cortex, whereas in younger 5xFAD mice, plaques were more abundant in the hippocampus than in the cortex, particularly for Aβ₄₂. The histological staining, where APP_sl_ mice showed a significant number of large plaques in the cortex, confirmed this hypothesis. Taken together, these findings suggest a direct link between brain amyloid pathology and salivary Aβ₄₂ concentrations, although confirmation in longitudinal studies using comparable models will be necessary. The high accuracy of our method and the correlation with brain pathology position the method as a potential tool for diagnostic screening, a matter of increasing importance concerning AD.

Building on these findings, the translational potential of the TIRS platform is considerable. With a sample requirement of only 5 µL and rapid readout, it could support wide-scale screening of at-risk populations, repeated monitoring in clinical trials, and biomarker-guided initiation of emerging amyloid-targeting therapies.(45) Beyond amyloid, the modularity of the analysis platform enables extension to p-tau and other neurodegenerative biomarkers, offering a flexible pipeline for multiplexed saliva-based diagnostics. Future studies will therefore prioritize three directions. First, validation in human cohorts will be essential to establish robustness across diverse populations, comorbidities, and disease stages. Second, longitudinal sampling in animals and patients will test whether salivary Aβ₄₂ dynamics reflect disease progression and therapeutic response. Third, expanding the biomarker panel to include p-tau isoforms, APP derivatives, and inflammatory mediators will enable multiplexed profiling. Such multidimensional datasets may benefit from advanced machine-learning approaches, where support vector machines and other classifiers could provide additional discriminative power beyond the single-marker setting explored here. Together, these steps define a clear translational path from proof-of-concept in mice toward clinically viable human diagnostics. By coupling ultrasensitive detection with minimal sample requirements, the TIRS platform addresses a central barrier in neurodegenerative diagnostics: enabling scalable, non-invasive, and repeatable measurements of disease-relevant proteins. More broadly, this work highlights saliva as an underexplored diagnostic resource. By uniting nanophotonic sensing with machine-learning analytics, the TIRS platform lays the foundation for affordable, accessible, and precision-guided diagnostics that could transform how neurodegenerative diseases are detected and monitored, shifting the paradigm from invasive, resource-intensive testing towards real-time, widely accessible, and population-scale screening.

## Materials and Methods

### Animal Models

Two AD mouse models of amyloidosis were used to detect salivary Aβ: 5xFAD and human APP_sl_ (APP_sl_) mouse models. Both show strong age-dependent plaque burden, making them ideal to test whether salivary Aβ reflects brain pathology.(46,47). By investigating two different models of AD, we want to prove that salivary Aβ is not an isolated event confined to one model but rather connected to the brain pathology in general. APP_sl_ mice overexpress human APP751_sl_ under the control of the Thy1 promoter with London (V717I) and Swedish (K670N, M671L) mutations, causing elevated Aβ₄₀ and Aβ₄₂ levels, and plaque formation in the brain;(48) cognitive deficits appear after 6 months.(49) The cohort included 18 female transgenic mice and 15 female non-transgenic littermates aged 16-17 months. The 5xFAD mouse model (The Jackson Laboratory, strain #008730) has been used in about 10% of studies (AlzPED(50)). The model is based on a C57BL/6J background and overexpresses human amyloid beta amyloid precursor protein due to the Swedish (K670N, M671L), Florida (I716V), and London (V717I) familial Alzheimer’s disease (FAD) mutations as well as the human presenilin 1 (PS1) protein by harboring two FAD mutations, M146L and L286V.(46) Both transgenes are modulated by the mouse Thy1 promoter to drive overexpression specifically in brain neurons.(48) The 5xFAD model (C57BL/6J background) carries three APP mutations and two PS1 mutations, all under the Thy1 promoter, with plaques visible by 2 months. We studied 20 transgenic (11 female, 9 male) and 14 wild-type littermates (9 female, 5 male), aged 8.5–9.5 months. Animals were provided by Scantox Neuro GmbH and housed in a 12-hour light, 12-hour dark cycle in the core facility for animal husbandry and breeding (CFL) at the Medical University of Vienna. APPsl mice were housed in groups of 5-6 animals. 5xFAD mice were housed in groups of 2-5 animals, depending on sex. Animal experiments were approved by the Austrian authorities under the animal experimentation license GZ 2024-0.044.300 and performed according to ethical guidelines by qualified personnel.

### Saliva Sampling

Saliva was collected using an adapted protocol presented in Zubeidat et al..(51) Using a sterile needle, a 0.5 mL sterile reaction tube (Biozym) was punctured in the lower part and then inserted into a 2 mL sterile reaction tube (Biozym). Cotton swabs were gently rotated in the mouth mouse, then centrifuged at 8000g for 3 minutes to recover the saliva in the 2 mL tube.(47) Yields varied between animals, with a range from 1-13 µL. Samples were frozen on dry ice immediately after centrifugation and kept in a –80°C fridge until analysis. Saliva samples were taken on multiple dates and time points over the timespan of one month. A precise list of sampling dates, times and acquired volumes is provided in the supplementary material (Supplementary Table 3). Contaminated with blood or dirt particles or insufficient (<5 µL) samples were excluded.

### TIRS-based detection system

Biomarkers were measured with a bench-top total internal reflection scattering (TIRS) microscopy setup optimized for nanoparticle scattering detection (Supplementary Figure S1). The system exploits a glass micro-prism optically coupled to a coverslip via immersion oil, generating an evanescent wave (EW) at the glass-saliva interface when illuminated by a 640 nm diode laser (10 mW), collimated through a 20 mm lens.(34) Elastic scattered light from nanoparticles was collected by a Nikon CSI Plan Fluor 40x objective (NA 0.6, WD 2.8-3.6 mm) and directed through a 200 mm tube lens to a CCD camera.

### Synthesis of functionalized nanoparticles for the selective recognition of AD biomarkers

Functionalized gold nanoparticles (NPs) of 15 nm in diameter were synthesized via the Turkevich synthesis and subsequently functionalized through a ligand exchange process to replace citrate with bifunctional polyethylene glycol (PEG) molecules.(52) Briefly, 1.5 mL of 30 mM HAuCl_4_· 3H_2_O was added to 48.5 mL of boiling water (stirring at 400 rpm, 250°C). The solution was stirred under heating for 15 minutes and then stirred without heating for an additional 30 minutes. Later, an exchange ligand process was implemented immediately after synthesis to functionalize the gold nano-surface with thiolated-PEG molecules.(36–38) In particular, an aqueous solution of 10 μM SH-PEG-COOH was added to 2 nM NPs in a 1:1 volume ratio and stirred at room temperature for at least 2 hours to reach [PEG]/[NPs] molar ratio of 5000. The concentration of NPs was determined according to the Lambert-Beer law, Eq. 1:

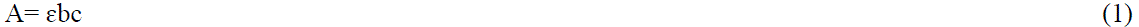

where:

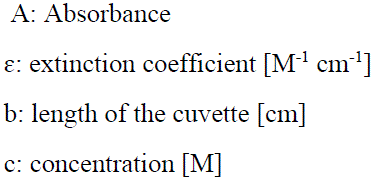

where ɛ is equal to 1.8 x 10^8^ M^-1^ cm^-1^.(53) Carboxyl groups on PEG were then exploited to covalently attach anti-β-amyloid 1-16 antibodies (Ab), targeting the N-terminal residues of the peptide.(37,39,54) Antibodies-bioconjugates were prepared starting from a 5 nM solution of the previously PEG-functionalized nanoparticles. Incubation was performed in 0.1 M PBS at a pH of 7.4 for 12 h at room temperature after activation with 0.4 mM EDC and 0.1 mM NHS (final concentration) for 30 min. A 1 mg/mL stock solution of antibody was added to 5 nM PEG-NPs in a 1:100 volume ratio. Samples were then centrifuged for 15 min at 4 °C and 10 000 rpm, and the pellet was redispersed in PBS 1x buffer to obtain 1 nM NPs-Ab. These NPs-Ab conjugates were characterized using standard techniques for colloidal systems. The UV-Vis absorbance spectrum exhibited a plasmonic peak at approximately 524 nm (Supplementary Figure S3a), consistent with 15 ± 1 nm diameter particles, as confirmed by dynamic light scattering (DLS, Figure S3b) and transmission electron microscopy (TEM), which also revealed uniform spherical morphology and good dispersion (Figure S3c).

### Saliva Concentration Measurements

As reported in Figure S4, 30 μL of capturing antibodies solution (23 μg/mL in PBS 1x) was sandwiched between a glass microscope slide and a glass coverslip to form a flow chamber and incubated for 15 minutes at room temperature, followed by three washes with PBS 1x. Saliva samples collected as described above were diluted 1:1 volume ratio with PBS 1x. Then, 10 μL of the diluted sample was flushed into the chamber and incubated for 5 minutes. Finally, 30 μL of nanoparticle–antibody conjugates (NPs-Ab, 1 nM) were introduced, incubated for an additional 5 minutes, and washed three times with PBS. The prepared chambers were placed on the sample stage of the TIRS-based microscope for scattering measurements.

### Image Processing and Analysis

For each sample, 30 images were acquired and analyzed using a Python script to quantify nanoparticle density (NPs/mm^2^), as reported in Supplementary Figure S8. The density of NPs in each image was calculated by dividing the number of detected particles by the image area (110 × 83 μm²). From these values, the mean NPs density and its corresponding standard error were determined.

### Tissue Sampling

For histological and biochemical analysis, 8 5xFAD mice (3 tg female, 4 tg male, 1 ntg female) aged 8.5-9.5 months and 16 APP_sl_ mice aged 17 months were sacrificed using an intraperitoneal pentobarbital injection. Mice were perfused with 0.9% NaCl solution and brains were sampled. The brain was divided into right and left hemispheres. The right hemisphere was fixed in freshly prepared 4% paraformaldehyde (Roth) in phosphate buffer (PBS) (PAN Biotech) for two hours at room temperature. Afterwards, the tissue was stored in a 15% sucrose (Sigma-Aldrich) in PBS (PAN Biotech) solution at 4°C overnight for about 12 hours of cryoprotection. After 12 hours right hemispheres were embedded into optimal cutting temperature (OCT) medium (Scigen) and frozen. The left hemisphere of the brain was dissected, cortex and hippocampus were isolated, and shock frozen on dry ice for biochemical analysis.

### Biochemistry

Cortex and hippocampus samples were tested for Aβ₄₂ and Aβ₄₀ concentration by using a ready-to-use ELISA kit for each of the proteins. (Mouse Aβ₄₂ ELISA Kit, KMB3441, ThermoFisher) (Mouse Aβ₄₀ ELISA Kit, KMB3481, Thermo Fisher) The Hippocampus mass was lysed via the addition of 150 µL of 5 M guanidine-HCl/50mM Tris, pH 8.0 buffer in 50 µL aliquots and the cortex mass was lysed using 1.2 mL of lysis buffer (5M guanidine-HCl/50mM Tris, pH 8.0) by 100 µL aliquot. Afterwards, the homogenates were incubated for 4 hours. Subsequently, homogenates were split and diluted in cold BSAT-PBS (1X PBS, 5% BSA, 0.03% Tween) for Aβ₄₀ ELISA or 1x PBS (Aβ₄₂) 1:5 and centrifuged at 16.000 x g for 20 min at 4°C. Pre-diluted samples were stored at –80°. For ELISA, samples were further diluted 1:250 (Aβ₄₂) and 1:100 (Aβ₄₀). Absorbance was measured using a microtiter plate reader CLARIOstarPlus, equipped with the SMART Control Software (BMG LABTECH, Servo LAB, Austria) at a wavelength of 450 nm.

### Histology

Right hemispheres embedded in OCT medium were put into formalin and then embedded into paraffin. Subsequently, the brains were cut coronally in a microtome and sections containing cortex and hippocampus were prepared for staining (Supplementary Figure S9). Aβ₄₂ and Aβ₄₀ were stained with Abeta 1-40 and 1-42 antibodies (both Antibodies Online, USA) according to a manual staining protocol. In brief, following deparaffinization and heat-induced antigen retrieval at pH6, tissue samples were incubated with rabbit polyclonal anti-Abeta 1-40 (clone AA 33-40, dilution: 1:1500) and 1-42 (clone AA 37-42, dilution 1:100) antibodies for 30 minutes, followed by a haematoxylin II counterstain for 20 minutes and a blue colouring reagent for 8 minutes. For detection, the universal anti-rabbit Dako EnVision+ System-HRP Labelled Polymer antibody (Dako, Agilent Technologies, USA) was incubated for 30 minutes. All stained slides were digitally imaged at 40X magnification using a Leica Aperio GT450 Dx digital slide scanner (Leica Biosystems, Chicago, USA).

### Statistical analysis

Salivary biomarker concentrations (Aβ₄₂, Aβ₄₀) were pre-processed to remove incomplete or erroneous observations. In total, 33 APP_sl_ mice and 34 5xFAD mice were sampled. For animals in which the collected saliva volume was sufficient, both Aβ₄₂ and Aβ₄₀ were analyzed; otherwise, one biomarker was prioritized to ensure adequate sample sizes for statistical analysis. Specifically, in APP_sl_ mice, n= 26 were analyzed for Aβ₄₂ and n= 14 for Aβ₄₀; in 5xFAD, n=26 for Aβ₄₂, and n=22 for Aβ₄₀. Because some animals had multiple saliva determinations on different dates, repeated measures were aggregated to a single subject-level value. Since concentration values showed outliers and non-normal distributions, repeated measures within animals were summarized by the within-subject median concentration (computed separately for each biomarker). Concentration values were then extracted for each group and biomarker, and all subsequent analyses were performed on these subject-level values. The following analyses were performed: ordered histograms to visualize the distribution of values in the two groups; boxplots with Student’s t-test to assess significant differences between group means. Group comparisons were performed using an unpaired two-sample t-test under the assumption of equal variances. Statistical significance was considered for p-values less than 0.05. A p-value above this threshold indicates that the observed difference is likely due to chance, and the null hypothesis of equal population means is not rejected. To evaluate the discriminative ability of each biomarker, receiver operating characteristic (ROC) curves were generated by plotting the True Positive Rate (TPR) against the False Positive Rate (FPR) across a range of thresholds. The area under the curve (AUC) was calculated to quantify the overall classification performance. Additionally, the Youden index was computed to identify the optimal threshold that maximizes the sum of sensitivity and specificity. For binary classification performance, the accuracy was computed as Eq. 2:

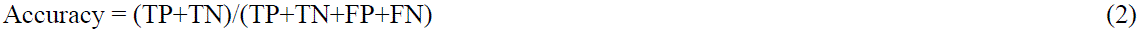

where:

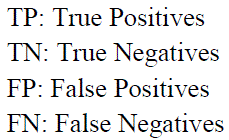

### Correlation Between Salivary and Brain Aβ₄₂ Levels in the APP_sl_ Model

For APP_sl_, n = 15 animals were available: for 5xFAD, n = 23 for the hippocampus and n = 24 for the cortex. Both Pearson’s correlation coefficient (r) and Spearman’s rank correlation coefficient (ρ) were calculated, as Pearson measures the strength and direction of a linear relationship between two continuous variables and ranges from −1 to +1, whereas Spearman is a nonparametric measure suitable for assessing monotonic relationships, particularly in data that are non-normally distributed or contain outliers. We first computed pooled correlations and then stratified the analysis by genotype to assess within-group associations. For each pair of variables, the correlation coefficient and its associated p-value were reported. A p-value below 0.05 was considered significant.

### Models: Logistic Regression and SVM

Salivary Aβ₄₂ was evaluated for classifying wild type (wt) vs. transgenic (tg) status using Logistic Regression and a Support Vector Machine (SVM, implemented as a Support Vector Classifier, SVC).(55) In both approaches, salivary Aβ₄₂ concentration was used as the single predictor of transgenic status (tg vs wt). Model performance was assessed with Leave-One-Out Cross-Validation (LOOCV). At each LOOCV iteration, one subject was held out for testing, while the remaining subjects were used for training. The model was then trained on the standardized training data and used to generate a prediction for the test subject. Repeating this across all folds yielded out-of-sample predictions for every subject. From the aggregated out-of-fold predictions, we computed the ROC curve, the area under the curve (AUC), the Youden-optimal decision threshold and confusion-matrix-based metrics. Uncertainty was estimated with 1000 non-parametric bootstrap resamples, where subjects were resampled with replacement, and the entire LOOCV pipeline was rerun. For Logistic Regression, this included scaling and refitting within each fold. For SVM, kernel and hyperparameters were selected once per bootstrap replicate via an inner crossvalidation on the bootstrap sample; the selected configuration was then refitted on each training fold and evaluated on the left-out subject within that replicate’s LOOCV. Across bootstrap replicates, we recorded AUC and confusion-matrix metrics and obtained 95% confidence intervals (CI) via the percentile method. For both logistic regression and SVM, we additionally derived CI for the optimal decision threshold, expressed in terms of NPs density.

## List of Supplementary Materials

Materials

Figs S1 to S9

Tables S1 to S4

## Acknowledgments

Authors would like to acknowledge the “Centro di competenza—RISE” funded by FAS Regione Toscana.

“I-PHOQS” project financed by the EU next generation PNRR action (CUP B53C22001750006)

“SENSOR project” (Agyr 2024 – Airalzh Foundation)

ERC Proof of Concept grant 101069344 OPTIMEYEZ

FFG grant 900435

“OPTO-19FLUIDIC” (CUP B53D23015530006) PRIN project financed by the EU next generation PNRR action

“DoptoScreen project” (Fondo di Beneficenza Intesa San Paolo 2019, B/2019/0289) Advance Lightsheet Microscopy Italian Mode of Euro-bioimaging ERIC

## Author contributions

Conceptualization: CD, MK, GL, FFC; Methodology: CD, GL, MK, LuP, LaP; Investigation: CD, GL, MK, FFC, LuP, LaP; Visualization: All authors; Funding acquisition: CD, CC, BB, FSP; Project administration: CD, GL, MK; Supervision: CC, AW, BB, RL, FSP; Writing – original draft: CD, GL, MK, FFC; Writing – review & editing: all authors

## Data and materials availability

All data are available in the main text or the supplementary materials

