## Supplemental File for "Ultrasensitive saliva-based detection of early Alzheimer’s disease biomarkers via nanoparticle-enhanced evanescent scattering microscopy"

Supporting Information file

*Materials*

Gold(III) chloride trihydrate (HAuCl_4_ ∙ 3H_2_O); trisodium citrate dihydrate (C_6_H_5_O_7_Na_3_ ∙ 2H_2_O); hydrochloric acid (HCl); nitric acid (HNO_3_ – 70%); Poly(ethylene glycol) 2‐mercaptoethyl acetic acid (SH PEG‐COOH, Mn 7500); N‐(3‐Dimethylaminopropyl)‐N‐ethylcarbodiimide hydrochloride (EDC); N‐Hydroxysuccinimide (NHS); Anti-Amyloid Antibody, β 1-40; Anti-Beta-Amyloid 1-42 Antibody; Amyloid β Protein Fragment 1-42; Amyloid β Protein Fragment 1-40 were purchased from Merck. Anti-β-Amyloid, 1-16 was purchased from BioLegend. 0.5 mL and 2 mL sterile reaction tube (Biozym), sterile needles, cotton swabs, 0.9% NaCl physiological solution for animals perfusion, 2 mL crytubes (Biozym) for freezing of hippocampus and cortex, 4% paraformaldehyde (Roth) in phosphate buffer (PBS) (PAN Biotech) sucrose (Sigma-Aldrich) in PBS (PAN Biotech), OCT medium (Scigen), Mouse Aβ₄₂ ELISA Kit, KMB3441 (ThermoFisher), Mouse Aβ₄₀ ELISA Kit, KMB3481 (Thermo Fisher), 5 M guanidine-HCl/50mM Tris (pH 8.0 buffer), BSAT-PBS (1X PBS), 5% BSA, 0.03% Tween, paraffin, Abeta 1-40 and 1-42 antibodies (both Antibodies Online, USA), anti-rabbit Dako EnVision+ System-HRP Labelled Polymer antibody (Dako, Agilent Technologies, USA).

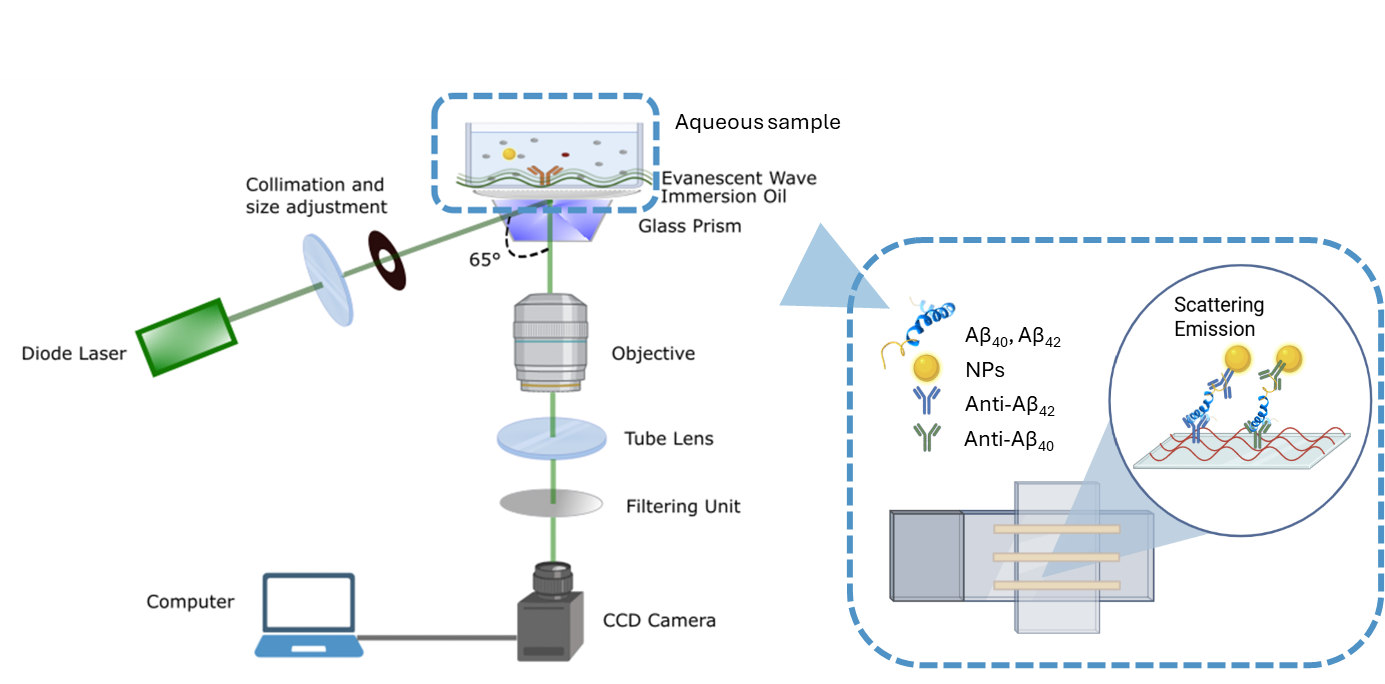

**Supplementary Figure 1**: **Scheme of the TIRS-based assay for detection of Aβ biomarkers**. A diode laser directed at a 65° incidence angle through a glass prism generates an evanescent wave at the sensing interface. Biological samples containing Aβ₄₀ or Aβ₄₂ peptides are incubated on a capture surface functionalized with specific antibodies. Gold nanoparticles (NPs) conjugated with detection antibodies bind to the immobilized peptides, forming a sandwich immunoassay. The interaction of the evanescent wave with NP–antibody complexes induce strong scattering emission, which is collected through an objective, filtered, and recorded by a CCD camera. The scattering intensity correlates with nanoparticle density at the surface, enabling quantitative measurement of Aβ₄₀ and Aβ₄₂ concentrations.

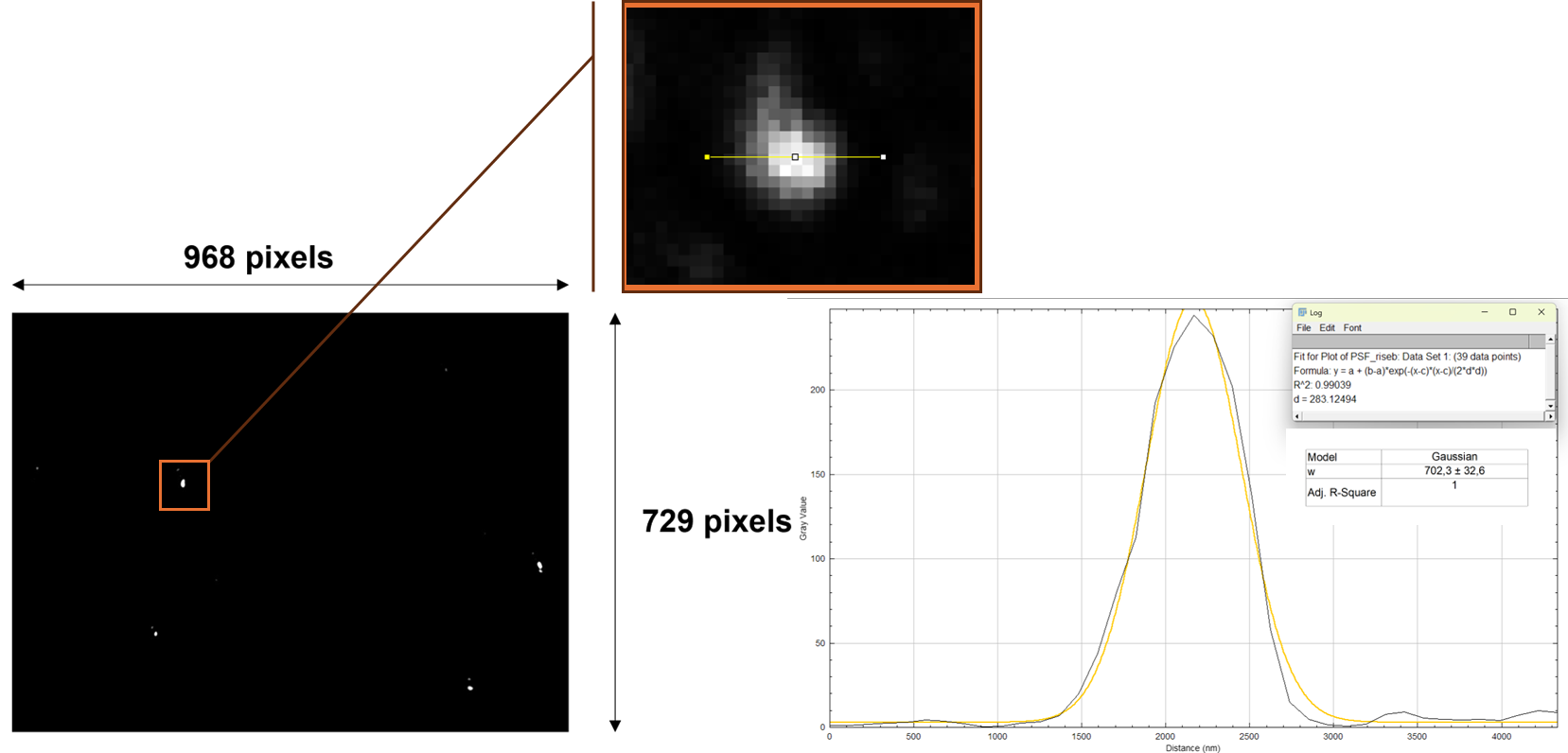

**Supplementary Figure 2**: Image acquired with the traditional TIR setup and zoom of the nanoparticle used to plot the PSF profile. PSF profile of the gold nanoparticle and its Gaussian fit. The FWHM (w value) of the Gaussian function provides the image lateral resolution.

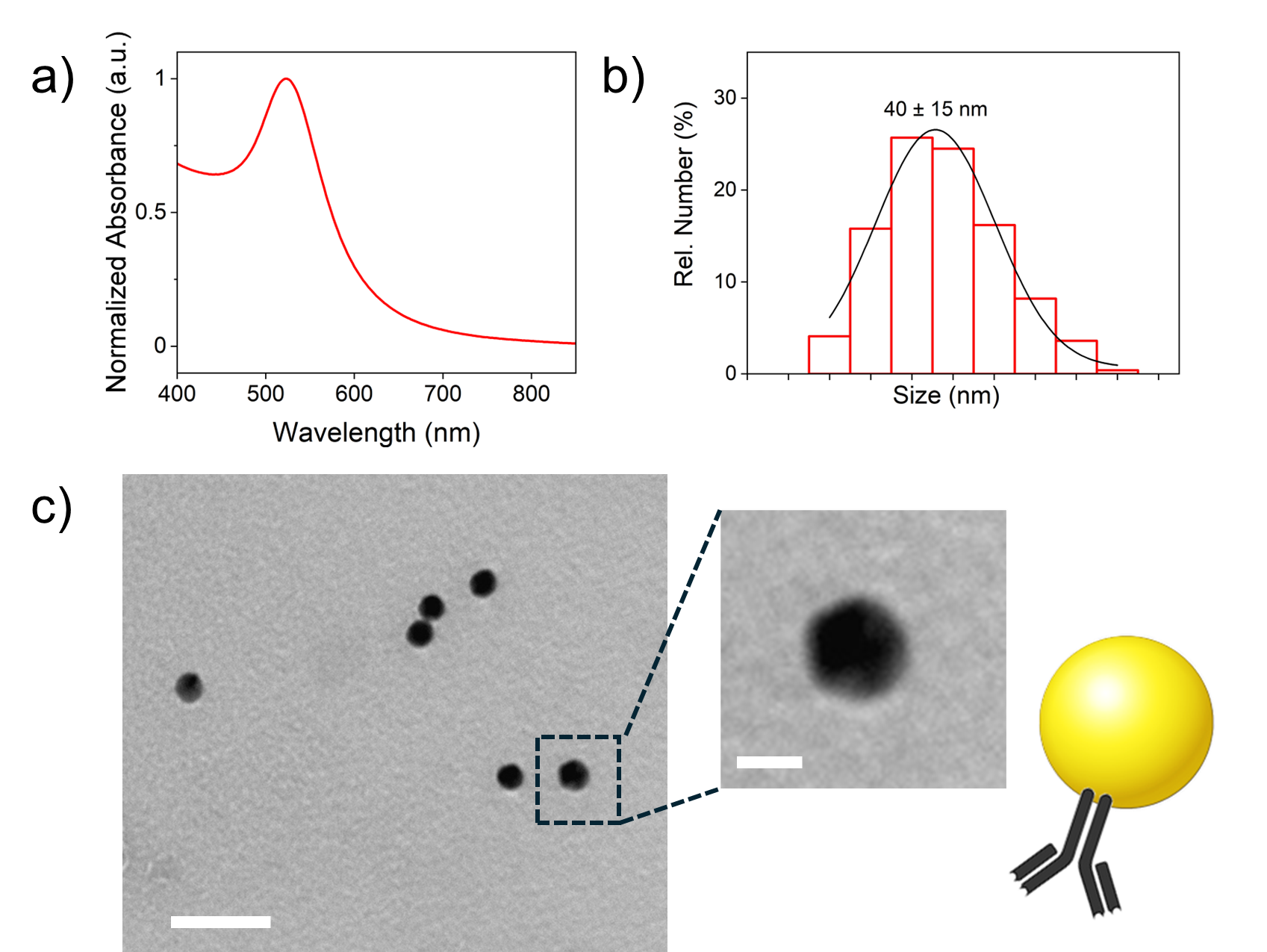

**Supplementary Figure 3**: Characterization of gold NPs covalently functionalized with anti-β-amyloid 1-16 antibodies in terms of plasmonic properties with UV-visible spectrum (a), hydrodynamic diameter via Dynamic Light Scattering (DLS) measurement (b), and morphology distribution with transmission electron microscopy (TEM) analysis (c). The scale bar is 50 nm for the TEM image in (c) and 10 nm for the inset.

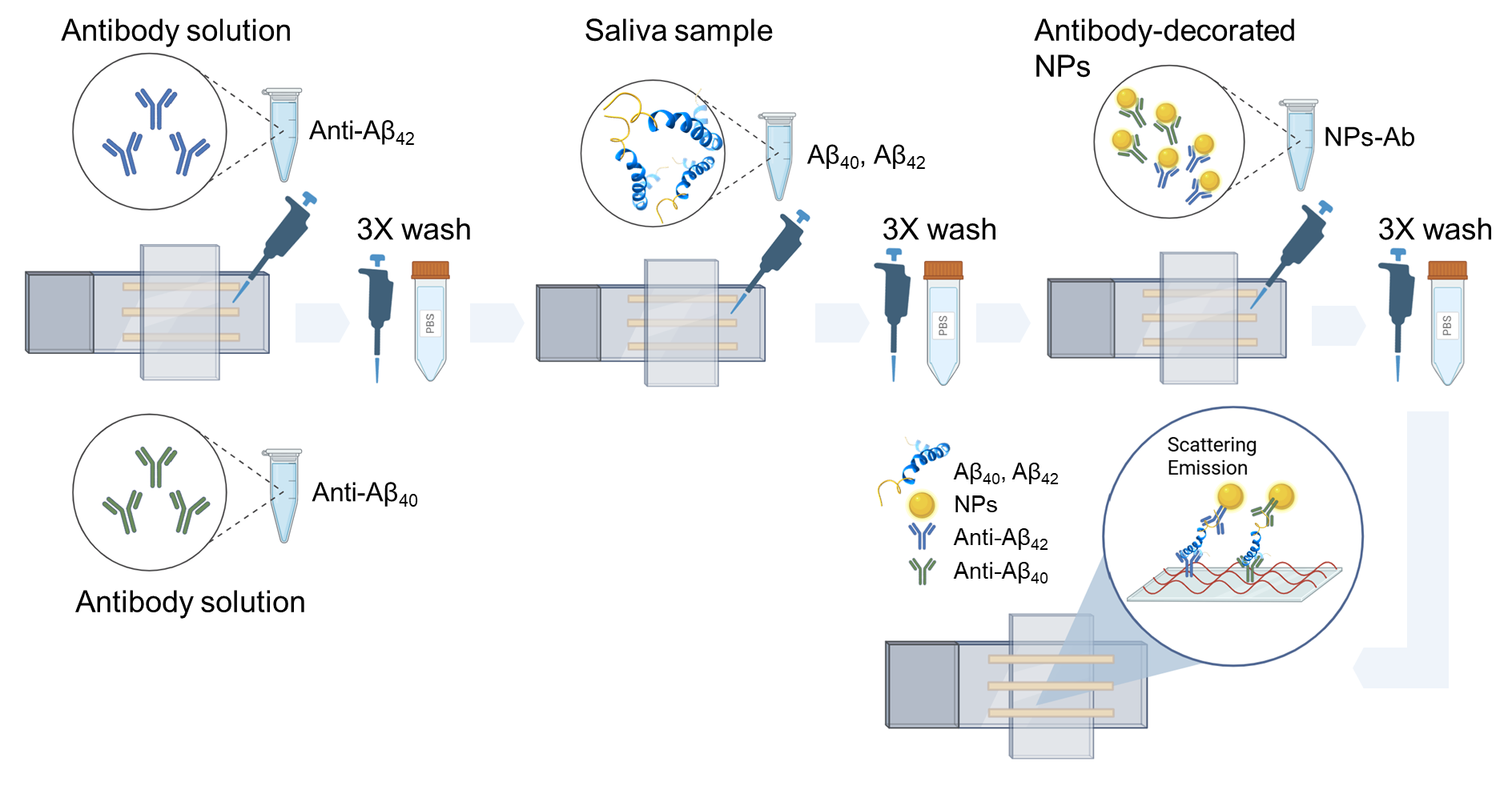

**Supplementary Figure 4**: **Schematic representation of the sample preparation steps.** After creating two channels on the microscope glass slide, the solutions are flushed into the channels in the order described in the figure, with an incubation time of 5 minutes at the first step and then 4 minutes for the subsequent steps.

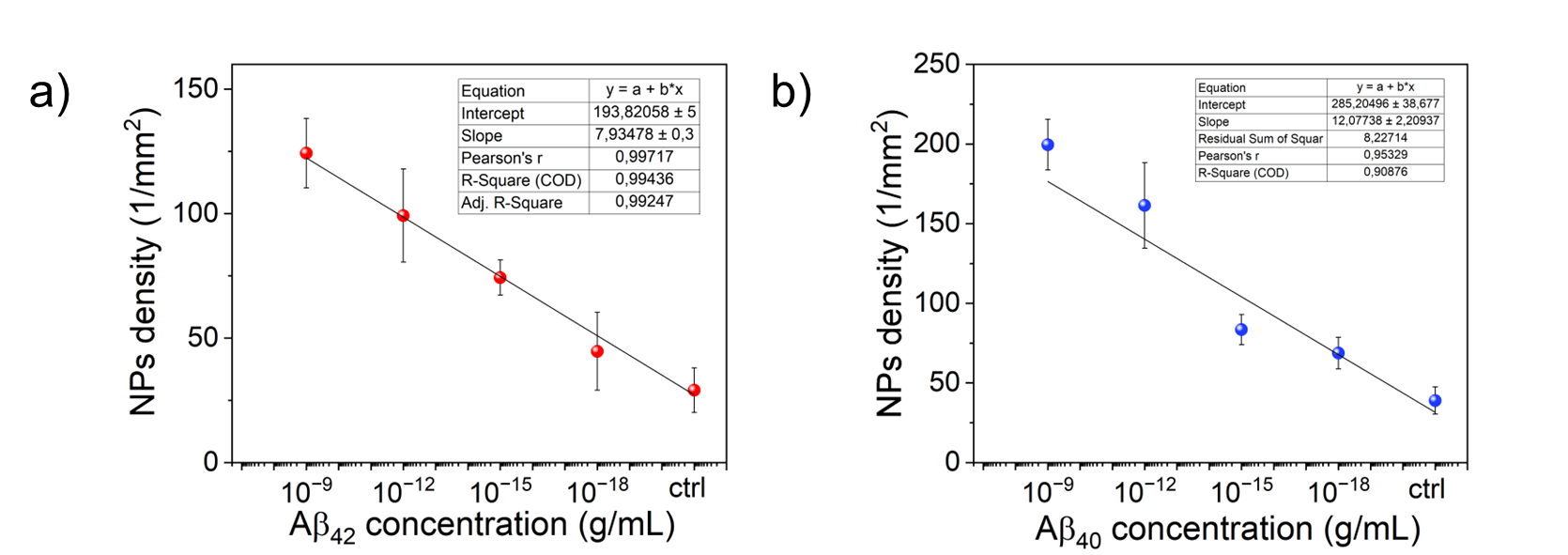

**Supplementary Figure 5:** Calibration curves for the scattering measurements performed with synthetic Aβ₄₂ (a) and Aβ₄₀ (b) concentrations ranging from 10^-9^ to 10^-18^ g/mL for each biomarker. The functions describing the calibration curves are:

Eq. (1): y [mm^-2^] = 193.8 + (7.9 log_10_ (x[g/mL]) ± 0.3

Eq. (2): y [mm^-2^] = 285.2 + (12.0 log_10_ (x[g/mL]) ± 2.2

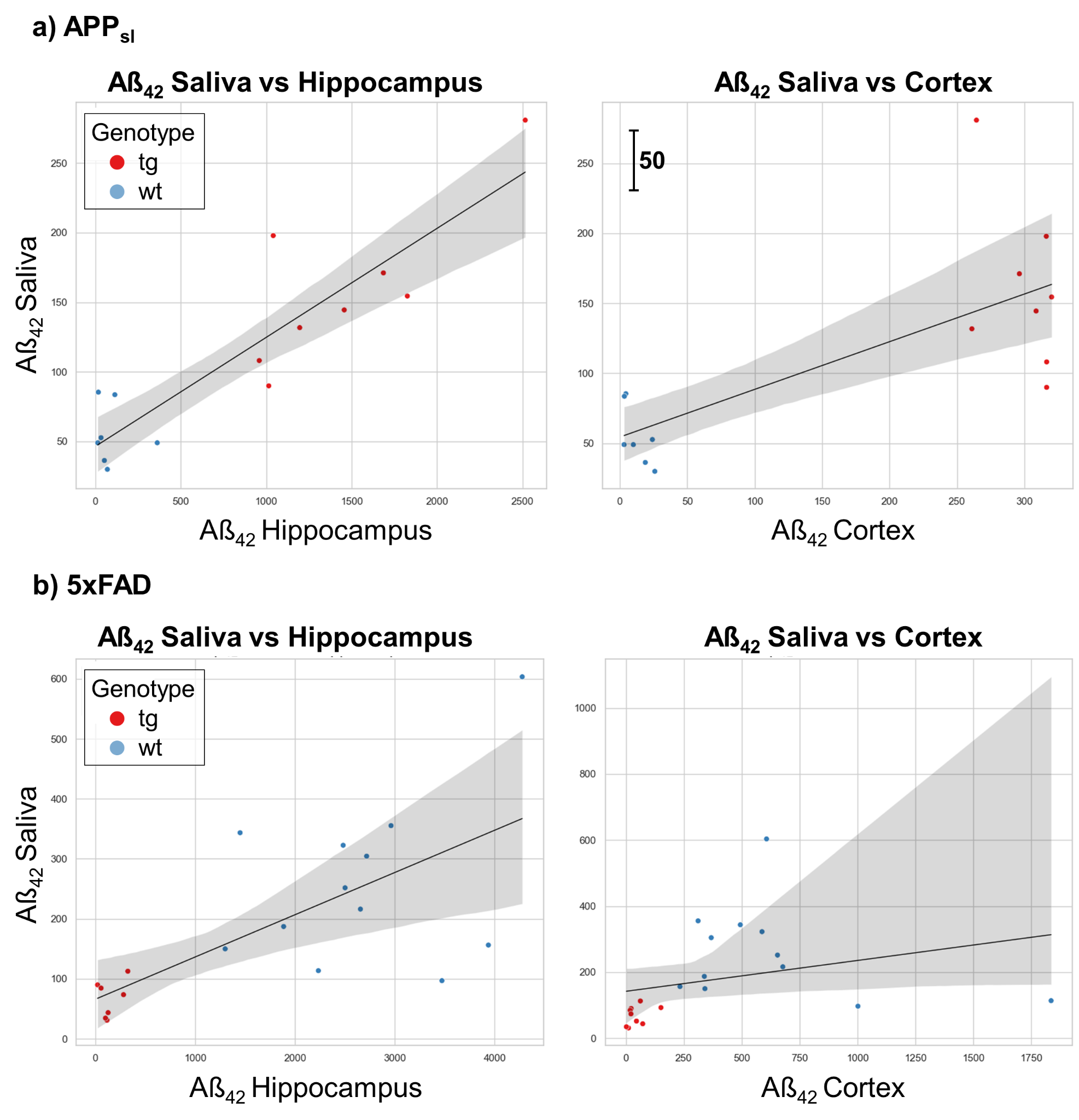

**Supplementary Figure 6**: Regression analysis to compare pooled Aβ₄₂ levels in saliva versus hippocampus (left panel) and in saliva versus cortex (right panel), for APP_sl_ (a) and 5xFAD (b). In the shaded area are the 95% confidence intervals of the regression line.

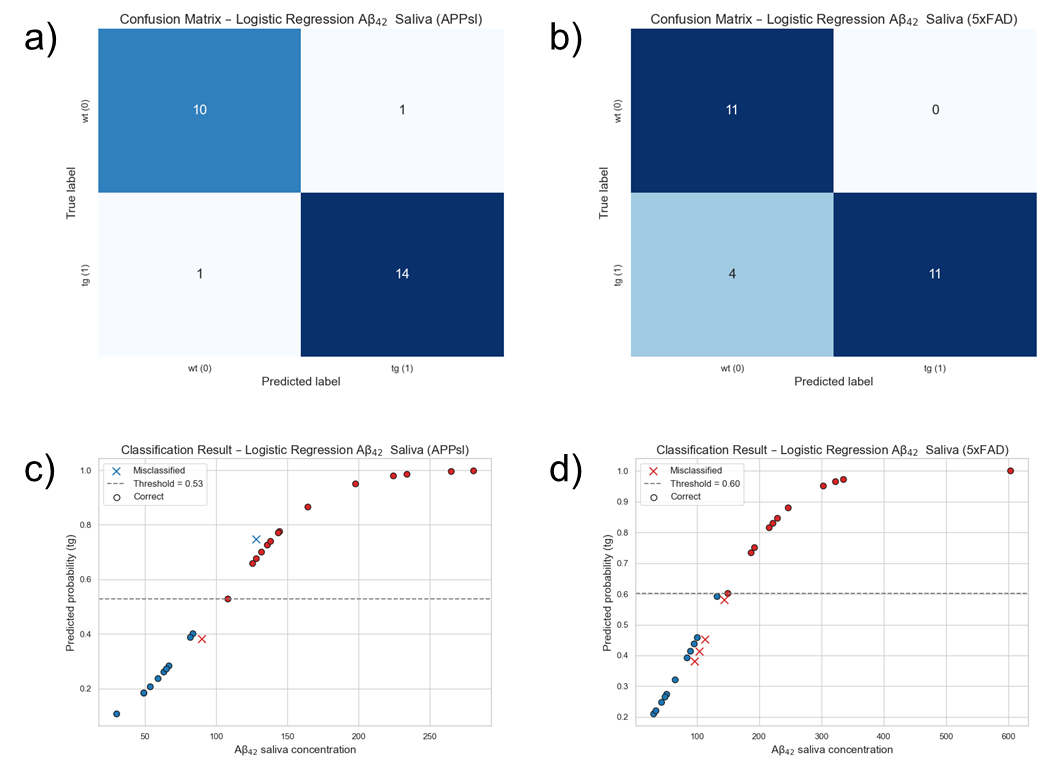

**Supplementary Figure 7:** Confusion matrices summarizing the performance of logistic regression in distinguishing wt from tg mice for both APP_sl_ and 5xFAD models based on salivary Aβ₄₂ concentrations. (a and b). Probability plots using the complete series of data (c and d).

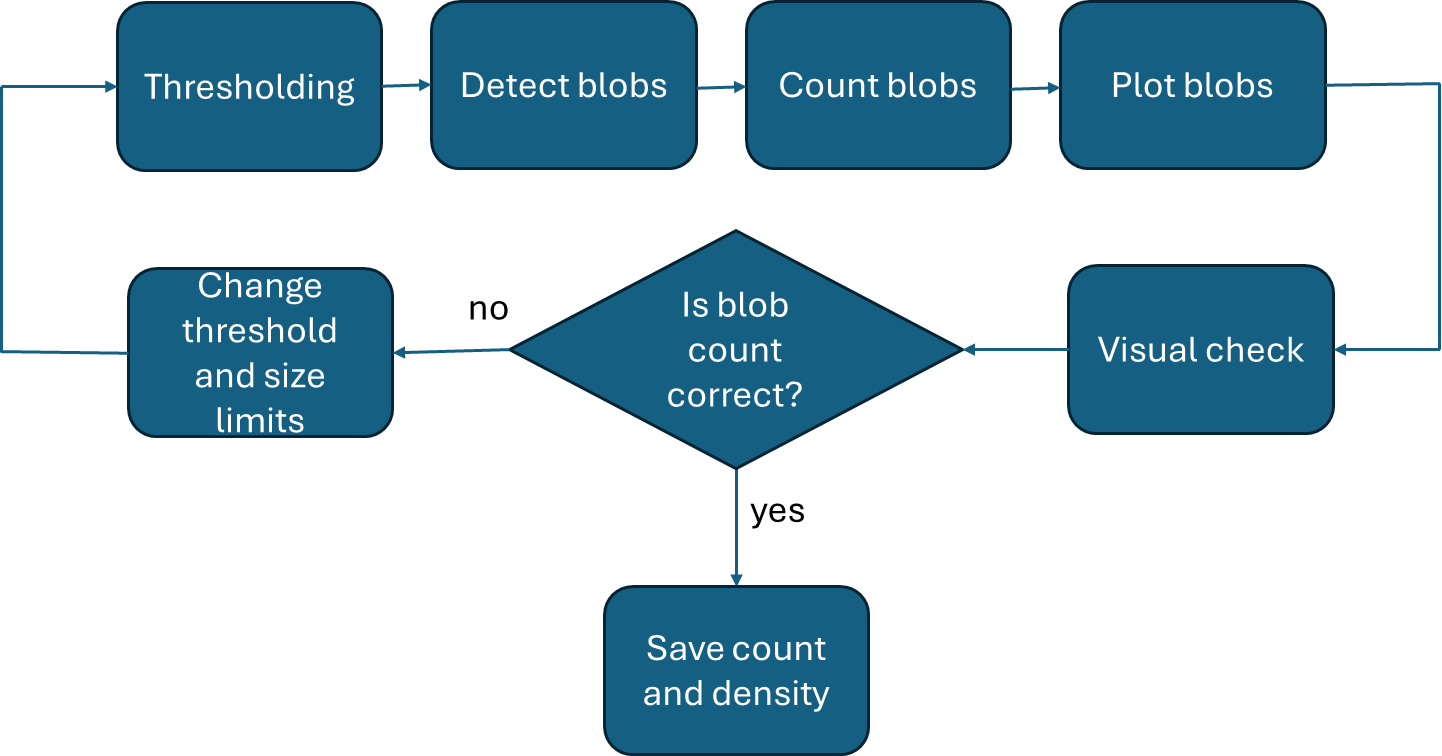

**Supplementary Figure 8**: **Workflow for detecting and quantifying particle density.** First, the image is binarized using a thresholding step. Next, a blob detection function identifies particles based on neighboring pixel values. Detected particles are then labeled, highlighted, and plotted for visual inspection. The semi-automated script enables the user to evaluate the quality of blob detection and, if necessary, adjust parameters such as the threshold level and minimum/maximum blob size before re-running the function to achieve satisfactory results.

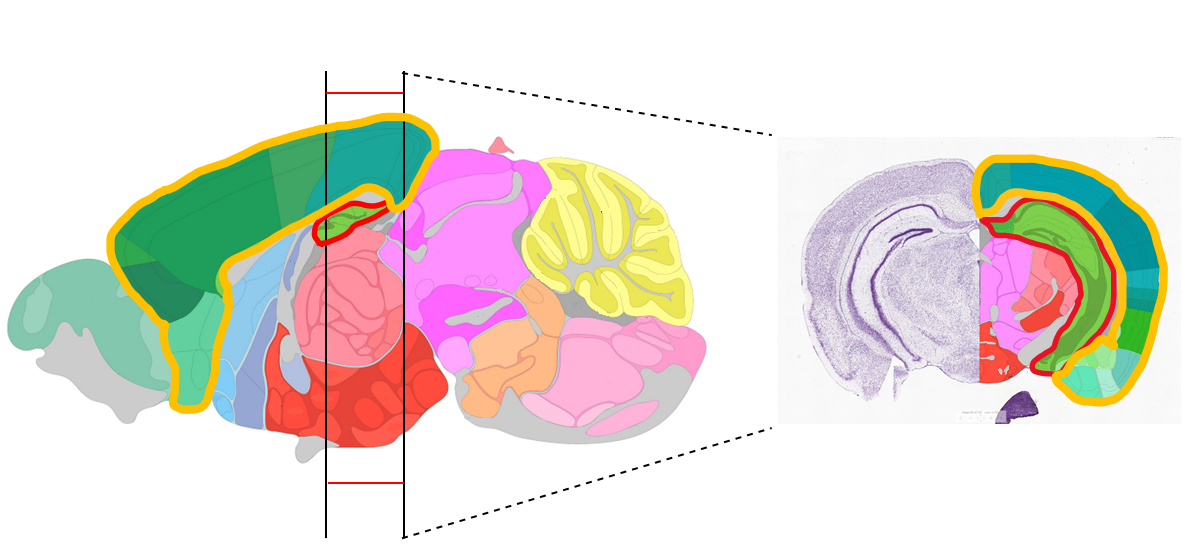

**Supplementary Figure 9**: Regions in the right hemispheres used for histological analysis. The brains were sagittally cut, and slides containing both hippocampus and cortex were used for staining. (Images taken from the Allen Brain Atlas) (https://mouse.brain-map.org/static/atlas)

*Supplementary Tables:*

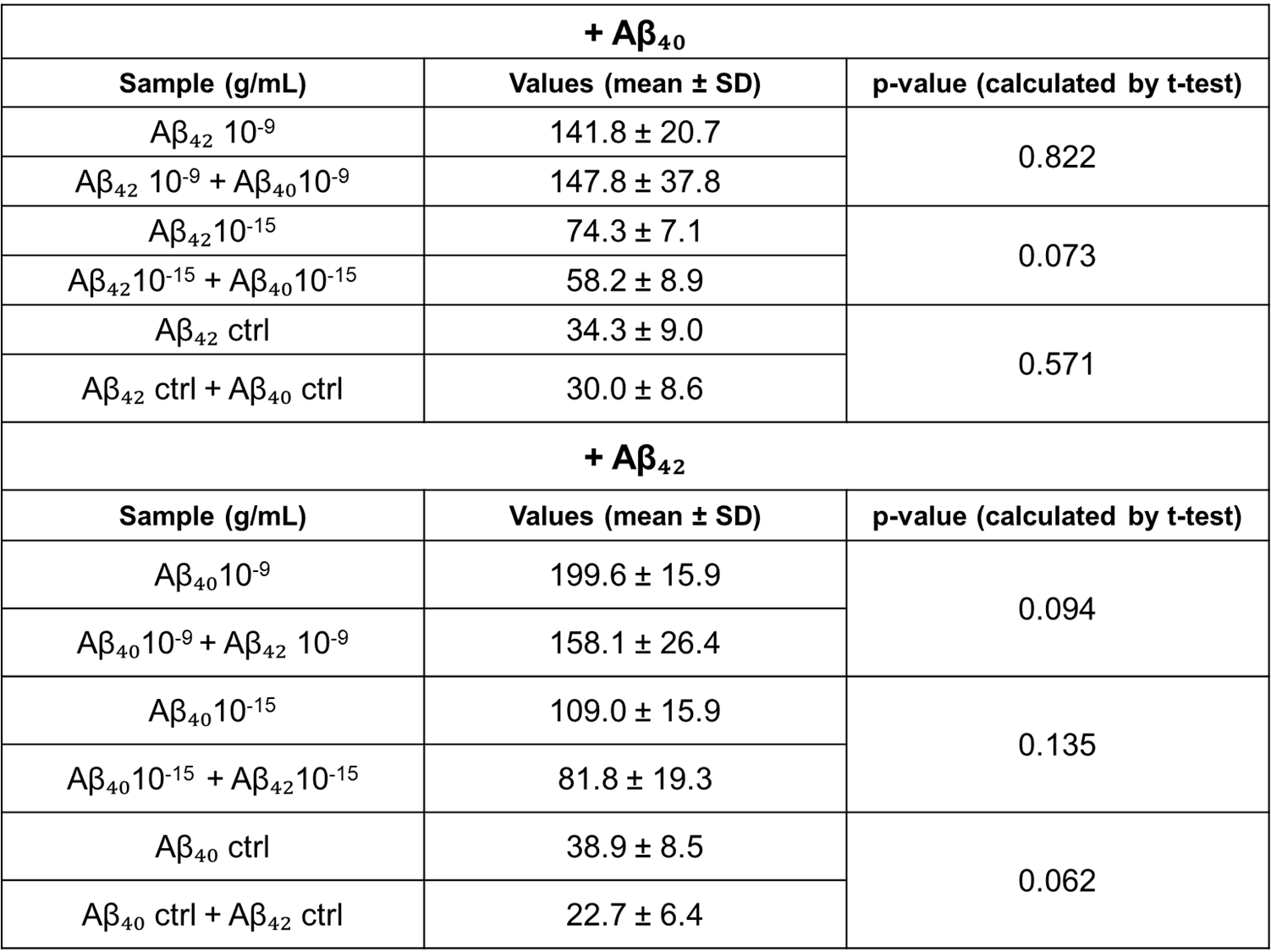

**Supplementary Table 1: Statistical evaluation of the specificity of the TIRS platform for Aβ₄₂ and Aβ₄₀ detection.** The table reports pairwise comparisons of nanoparticle density changes measured in solutions containing Aβ₄₂ alone, Aβ₄₀ alone, or both species. Non-significant p-values confirm the absence of cross-reactivity between the two amyloid species. In contrast, significant p-values indicate specific recognition of the target peptide across a concentration range of 10⁻¹⁸ g/mL and below.

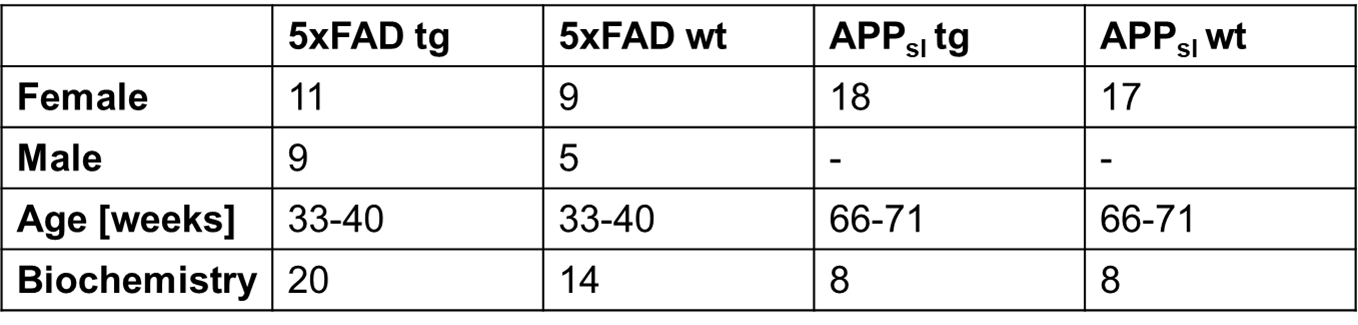

**Supplementary Table 2**: Number and age of tested animals with sex, genotype and number of animals used for biochemistry.

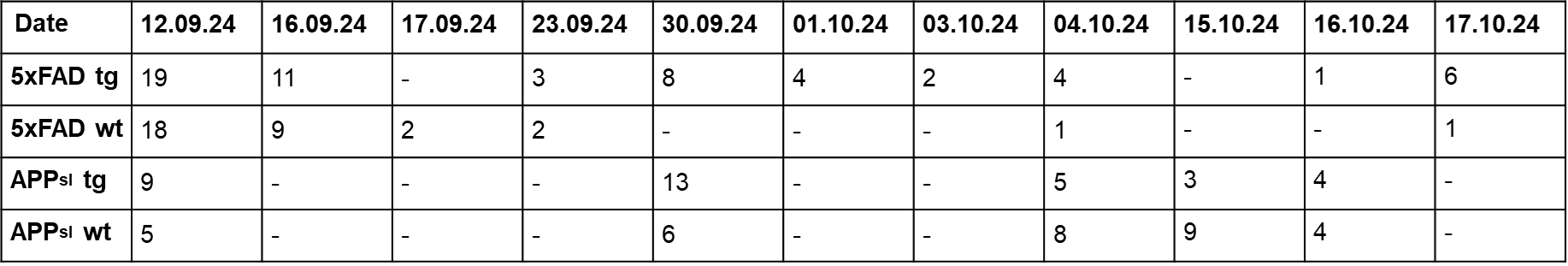

**Supplementary Table 3**: Number of analyzed samples per sampling date and strain/genotype.

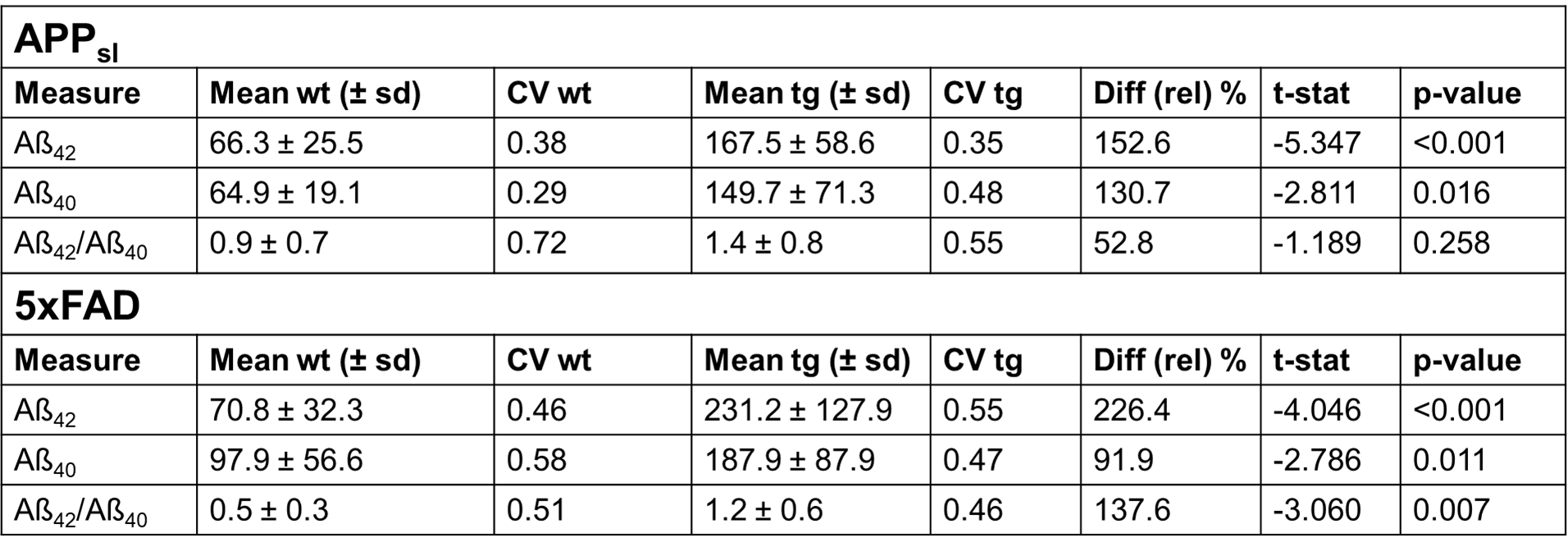

**Supplementary Table 4: Quantitative analysis of Aβ biomarkers in saliva samples from APP_sl_ and 5xFAD transgenic and wild-type mice.** The table reports mean concentrations, coefficients of variation (CV), relative differences, and p-values, for Aβ₄₂, Aβ₄₀, and the Aβ₄₂/Aβ₄₀ ratio in each model.
